# Data-driven machine learning for pattern recognition supports environmental quality prediction for irrigated rice in Brazil

**DOI:** 10.1101/2022.06.02.494614

**Authors:** Germano Costa-Neto, David Henriques da Matta, Igor Kuivjogi Fernandes, Luís Fernando Stone, Alexandre Bryan Heinemann

## Abstract

The sustainability of irrigated rice (*Oryza sativa L*.) production systems in Brazilian tropical region highly depends on the success of developing stable cultivars. To achieve this goal, many steps in product development must address the environmental variability and genotype by environment interactions (GE), which makes difficult the design and development of local-specific adapted cultivars. Thus, the adoption of new strategies for characterizing environmental-phenotype relations are the key for optimizing this process. In addition, it could also benefit post-breeding stages of seed production. To overcome this situation, we implemented a data-driven approach to link environmental characterization to yield clustering using historical data (1982-2017, 31 locations, 471 genotypes), 42 envirotyping covariables and machine learning (ML), combining two unsupervised (K-means and decision tree models, DTC) algorithms. Additionally, linear mixed models (LMM) were applied to explore the relations between the outcomes of our approach and GE analysis for irrigated rice yield in Brazilian tropical region. Four environments were identified: Very Low Yield (1.7 Mg.ha^-1^), Low Yield (5.1 Mg.ha^-1^), High Yield (7.2 Mg.ha^-1^), and Very High Yield (9.0 Mg.ha^-1^), considering all genotypes and regions. Our approach allows the prediction of environments (yield clusters) for a diverse set of growing conditions and revealed geographic and climatic causes of environmental quality, which differ according to each region and genotype group. From the LMM analysis, we found that the current relation between genetics (G), environmental variation (E), and GE for rainfed rice in Brazil is 1:6:2, but when we introduced our data-driven clusters (ME), the ratio decreased to 1:5:1. Consequently, the selection reliability for local adaptability across an extensive region increases. Our approach helps to identify mega-environments in Brazil that could be used as a target population of environments (TPE) of breeding programs. Additionally, it helps to identify more productive and stable seed production fields.

**Highlights:**

- A nationwide environmental characterization and its relation to the genotype by environment interaction (GE) for grain yield of rainfed rice growing regions in Brazil.
- A data-driven approach capable to identifying clusters of yield levels and a machine learning approach to relate those clusters with environmental typologies.
- Unrevealed geographic and climatic causes of environmental quality for a group of genotypes or cultivar-specific predictions.
- The strategy benefits diverse stages of breeding (multiple environmental trial analysis) and post-breeding (selection of fields for seed production) as an alternative approach to reduce costs and support decisions on cultivar planting locations.

## 1 INTRODUCTION

Brazil is the largest rice producer outside of Asia. Rice plays a strategic role in Brazilian economy and society (USDA, 2018). Two rice production systems predominate in Brazil: irrigated (IR) and upland rice (rainfed). IR accounts for 90% of the total production and for 78% of the cultivated area (CONAB, 2021). Therefore, IR performance is increasingly important to ensure food and nutrition safety. For that, IR depends on water security (ANA, 2020). IR is produced in two ecosystems: subtropical (south of Brazil) and tropical (Southeast, Midwest, North, and Northeast megaregions of Brazil).

The subtropical region accounts for 85% of the Brazilian IR production (2020/2021 crop season), with an average yield of 8.6 Mg ha^-1^. On the other hand, tropical IR accounts for only 14% of the national IR production and has an average yield ranging from 6.0 Mg ha^-1^ (Northeast megaregion), 6.1 Mg ha^-1^ (North megaregion) and 6.4 Mg ha^-1^ (Midwest megaregion) to 6.6 Mg ha^-1^ (Southeast megaregion) (CONAB, 2021). As rice is a major staple food in Brazil, in order to ensure food security and supply logistics for the Brazilian population, we cannot rely solely on rice produced in the subtropical ecosystem, which is located in the extreme south of the country. An example of this vulnerability is the deleterious effect of the “ENSO” phenomenon on subtropical rice yields in Rio Grande do Sul due to extreme events of rainfalls or drought (Cai et al., 2020; Grimm et al., 2020).

In terms of rice logistic in the tropical regions (Midwest, North, Southeast, and Northeast megaregions), we emphasize that the supply should be greater through an increase in crop yield and the consequent strengthening of the rice processing industry located in that region. In this context, rice supply diversification by improving yields of tropical IR is strategic for Brazil. This region is characterized as having a lower yield and a higher variability compared to subtropical irrigated rice. This is mainly due to the harmful effects of certain biotic and abiotic factors on the crop. As for biotic factors, high incidence of fungal diseases, such as blast (*Magnaporthe oryzae*), pest insects, such as stink bugs (*Oebalus spp*.), and weeds are widely spread; as abiotic factors, thermal stresses are the most expected, implicating in the shortening of the crop cycle and lower response to fertilizers, especially nitrogen (Meus et al., 2021).

Maximizing genetic gains in IR in a tropical ecosystem requires a better environment characterization that aims to produce information that may assist breeding strategies to develop adapted yield germplasm for a certain production region. Breeding activities for IR have focused basically on direct grain selection and wide adaptation to an undivided target environment (tropical and subtropical) (Heinemann et al., 2015). Nevertheless, in a large and heterogeneous TPE, genotypes may react differently to environmental variation, resulting in genotypic rank changes (Windhausen et al., 2012). Thus, a better understanding of target population environments (TPE) is essential for IR breeding efforts committed to developing and identifying improved genotypes that are in some way superior considering plant production purposes in a tropical region. TPE is defined as the set of environments, including spatial and temporal variability, to which improved crop varieties developed by a breeding program needs adaptation (Nyquist 1991; Cooper et al., 1997; Crespo-Herrera et al. 2021, Carcedo et al., 2022).

Information on the TPE explains and predicts a significant component of genotype × environment x management interactions (G × E x M) for grain yield, enabling the optimization of environmental screens and breeding itineraries (e.g., Chapman et al., 2000a, 2000b, 2000c; Crespo-Herrera et al., 2021). A classification of spatial-temporal variations in yield levels across environment groups, as well as a determination of prevalent environmental covariables and their capacity of discrimination (i.e., magnitude in yield changes) within those environment groups, may support breeding programs that aim to develop better genotypes adapted to tropical IR (Heinemann et al., 2008, 2015). Here we presented a data-driven approach for pattern recognition and grouping environments based on quality in the context of breeding trials.

Here we interpret “environmental quality” as the potential productivity level of a given environment (field trial, that is, a combination of location, planting date, and management). Our approach is implemented in three steps: (1) gathering and characterizing large-scale environmental information (envirotyping) and historical yield data from breeding trials; (2) pinpointing clusters of yield levels using a unsupervised learning approach and; (3) applying an unsupervised learning approach to associate envirotyping data and environmental clusters as found in step 2. Finally, a permutation scheme using nationwide data from the Rice Breeding program of the Brazilian Corporation of Agricultural Research (EMBRAPA) is used as proof-of-concept of the methodology. We were able to identify the climatic drivers of GE across diverse regions in Brazil, mostly the pattern related to genotype by location interactions and its repeatability across years. Implications of these results should help the design of TPEs and support post-breeding decisions, such as product targeting (cultivar recommendation), and optimizing seed production sites.

## 2 MATERIAL AND METHODS

### 2.1 Regional setting in Brazil, South America

The study area encompasses a wide range of edaphoclimatic conditions across several climatic zones in Brazil (Figure 1). The geographic area (latitude and longitude) ranges from 7º S to 29º S and from 36º W to 58º W, respectively, covering one of the most important rice production regions outside of Asia. This study considered four megaregions (North, NO; Midwest, MW; Southeast, SE; and Northeast, NE), which are in 15 Brazilian States (TO, GO, RR, RJ, PB, PI, CE, MS, PR, ES, SE, AL, AP, MA, and SP). This covers about 36 locations, here represented by their municipalities (Dueré, Goianira, Formoso Do Araguaia, Canta, Lagoa Da Confusão, Macaé, Campos Dos Goytacazes, Flores De Goiás, Sousa, Teresina, Buriti Dos Lopes, Iguatu, Dourados. Irati, Morretes, Vitoria, Luiz Alves, Aracaju, Rio De Janeiro, Penedo, Mazagão, Arari, Riacho Dos Cavalos, Barbalha, Lima Campos, Morada Nova, Linhares, Propriá, Miranda, Rio Brilhante, Pium, Guarantigueta, Pindamonhangaba, Tremembé, São Mateus, and Itapecuru Mirim). In this region, the most important and commonly used cultivars are “IRGA 424,” developed for subtropical ecosystems, with average yield of 6,700 kg ha^-1^ (Silva et al., 2021), and “BRS Catiana”, developed for tropical and subtropical conditions, with an average yield of 7,253 kg ha^-1^, according to Morais et al. (2016).

**Figure 1.**
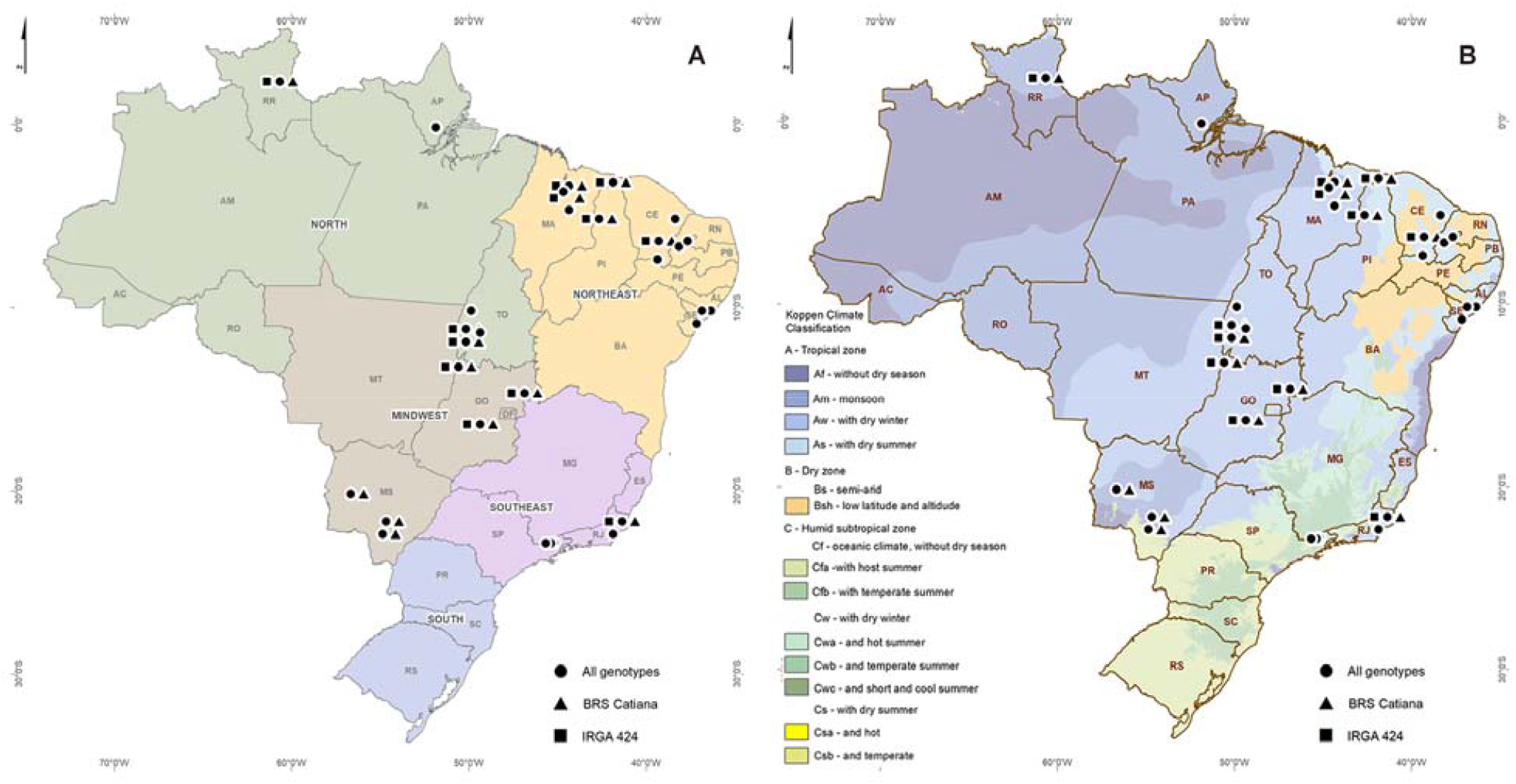
Geographic location of tropical irrigated rice trials in Brazil. (**A**) Geographic distribution of the macrorregions Midwest (MW). Southeast (SE), Northeast (NE) and North (NO). (**B**) Köppen climate classification. For both panels, black circle dots, black triangle and black square correspond to the distribution of the locations involving the genotypes groups “All genotypes” and the cultivar-specific “BRS Catiana” and “IRGA 424”, respectively.

### 2.2 Historical nationwide data set

This study uses a large, accumulated yield data set formed by variety trials on commonly grown and well-adapted irrigated rice varieties that derivate from the EMBRAPA Rice Breeding Dataset (Breseghello et al., 2021). As standardized by the EMBRAPA nationwide rice breeding program, each field trial comprises the best-performing 20 genotypes of the current elite germplasm. The experiment is conducted with randomized blocks and three replications. The trials comprise the years between 1982 and 2017, counting 21,749 phenotypic records, which mostly of the advanced yield trials considered for validating the methodology are comprised between 2011 and 2017 years. During this period, 471 elite lines were considered, among which two main cultivars are highlighted for the analysis below. However, in each year, some genotypes are discarded and others are included, thus the main cultivars (used as checks) and the best-performing lines always remain. On those trials, we selected only the trials for irrigated rice located at the tropical region (Figure 1). The following agronomic information was available for them: grain yield (GY, kg per ha), sowing date (SD), emergence date (ED), flowering date (FD, days after emergence - DAE), and maturation date (MD, DAE).

### 2.3 Genotype Groups

In order to conduct the analysis below, the full data set was split into three groups (hereafter called “genotype groups”). The first genotype group comprises 203 trials from 1982 to 2017 considering all genotypes tested across those years (Figure 1). Because of this, we hereafter refer to it as “All genotypes,” and this is the reference group of our study. Secondly, in order to verify the potential of the methodology to identify and predict environmental clusters, we created another two groups based on the two main cultivars adopted by growers in Brazil. The first was the “IRGA 424” group, named after the homonym cultivar IRGA 424. This set contains 47 trials from 2013 to 2017, of which we considered only the agronomic information for the cultivar IRGA 424. Likewise, the group “BRS Catiana” contains 57 trials conducted from 2006 to 2017, of which we considered the agronomic information only for the cultivar BRS Catiana.

### 2.4 Envirotyping protocol

Envirotyping (environmental + typing) is the process of using environmental information (e.g., weather data) to draw a profile of the growing conditions in terms of typologies affecting yield levels in a certain region. Here, this process was conducted using the software R (R Core Team. 2021) and following the steps of (1) collection of environmental data, (2) statistical processing of these data for the entire crop lifetime or for specific development stages, and (3) use of data for running phenotype-environmental associations in order to create “environmental clusters” (Table 1). For the latter, the agronomic variables (GY - grain yield, SD - sowing date, EMD - emergence date, FD - flowering date, and MD - physiological maturation date) were related to large-scale environmental information, such as daily climate data, considering “All genotypes” and the groups “IRGA 424” and “BRS Catiana.”

**Table 1.**
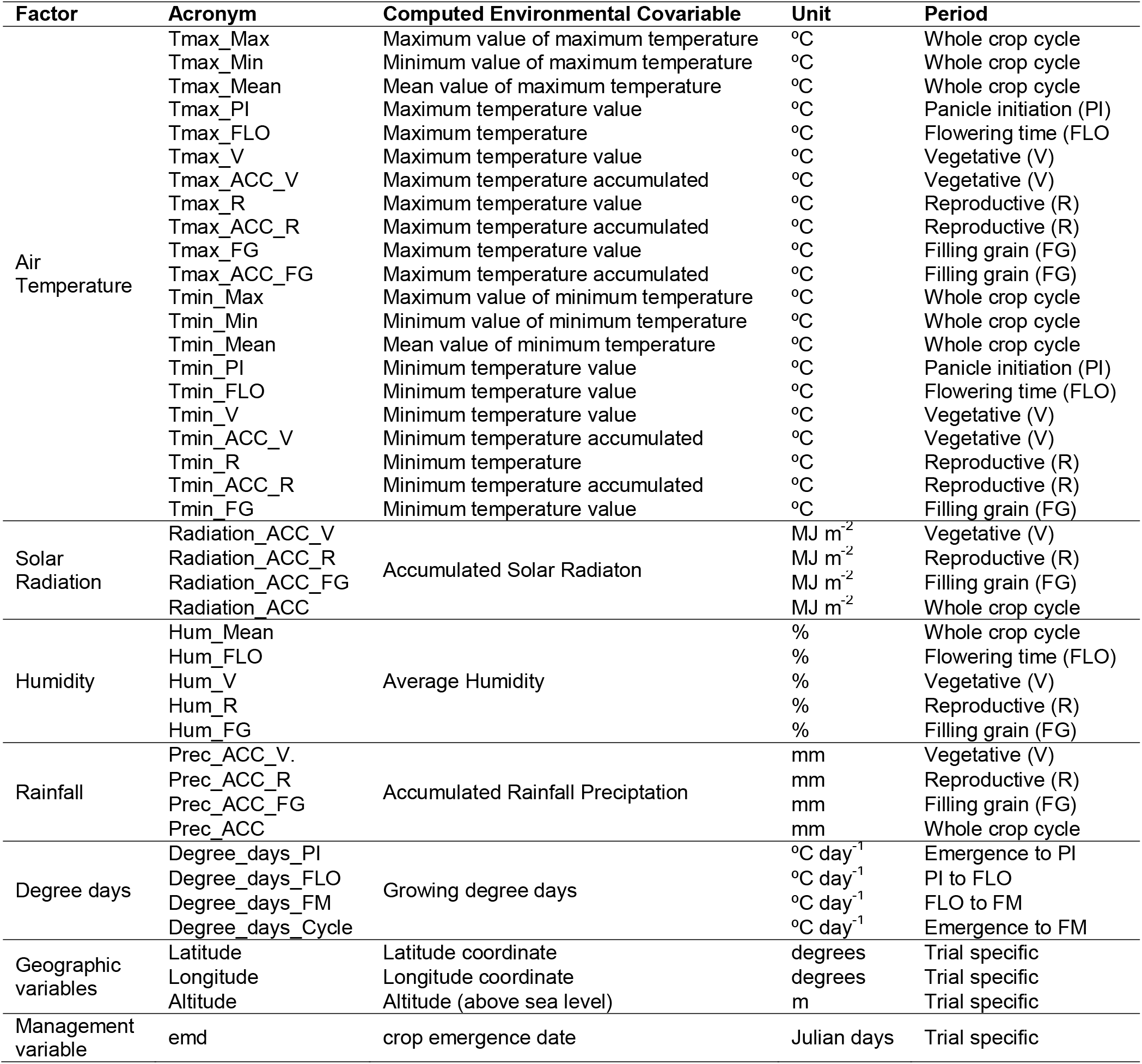
Acronyms for the environmental covariates (EC) applied in this study.

| Factor | Acronym | Computed Environmental Covariable | Unit | Period |
| --- | --- | --- | --- | --- |
| Air Temperature | Tmax_Max | Maximum value of maximum temperature | °C | Whole crop cycle |
|  | Tmax_Min | Minimum value of maximum temperature | °C | Whole crop cycle |
|  | Tmax_Mean | Mean value of maximum temperature | °C | Whole crop cycle |
|  | Tmax_PI | Maximum temperature value | °C | Panicle initiation (PI) |
|  | Tmax_FLO | Maximum temperature | °C | Flowering time (FLO) |
|  | Tmax_V | Maximum temperature value | °C | Vegetative (V) |
|  | Tmax_ACC_V | Maximum temperature accumulated | °C | Vegetative (V) |
|  | Tmax_R | Maximum temperature value | °C | Reproductive (R) |
|  | Tmax_ACC_R | Maximum temperature accumulated | °C | Reproductive (R) |
|  | Tmax_FG | Maximum temperature value | °C | Filling grain (FG) |
|  | Tmax_ACC_FG | Maximum temperature accumulated | °C | Filling grain (FG) |
|  | Tmin_Max | Maximum value of minimum temperature | °C | Whole crop cycle |
|  | Tmin_Min | Minimum value of minimum temperature | °C | Whole crop cycle |
|  | Tmin_Mean | Mean value of minimum temperature | °C | Whole crop cycle |
|  | Tmin_PI | Minimum temperature value | °C | Panicle initiation (PI) |
|  | Tmin_FLO | Minimum temperature value | °C | Flowering time (FLO) |
|  | Tmin_V | Minimum temperature value | °C | Vegetative (V) |
|  | Tmin_ACC_V | Minimum temperature accumulated | °C | Vegetative (V) |
|  | Tmin_R | Minimum temperature | °C | Reproductive (R) |
|  | Tmin_ACC_R | Minimum temperature accumulated | °C | Reproductive (R) |
|  | Tmin_FG | Minimum temperature value | °C | Filling grain (FG) |
| Solar Radiation | Radiation_ACC_V | Accumulated Solar Radiaton | MJ m<sup>-2</sup> | Vegetative (V) |
|  | Radiation_ACC_R |  | MJ m<sup>-2</sup> | Reproductive (R) |
|  | Radiation_ACC_FG |  | MJ m<sup>-2</sup> | Filling grain (FG) |
|  | Radiation_ACC |  | MJ m<sup>-2</sup> | Whole crop cycle |
| Humidity | Hum_Mean | Average Humidity | % | Whole crop cycle |
|  | Hum_FLO |  | % | Flowering time (FLO) |
|  | Hum_V |  | % | Vegetative (V) |
|  | Hum_R |  | % | Reproductive (R) |
|  | Hum_FG |  | % | Filling grain (FG) |
| Rainfall | Prec_ACC_V. | Accumulated Rainfall Precipitation | mm | Vegetative (V) |
|  | Prec_ACC_R |  | mm | Reproductive (R) |
|  | Prec_ACC_FG |  | mm | Filling grain (FG) |
|  | Prec_ACC |  | mm | Whole crop cycle |
| Degree days | Degree_days_PI | Growing degree days | °C day<sup>-1</sup> | Emergence to PI |
|  | Degree_days_FLO |  | °C day<sup>-1</sup> | PI to FLO |
|  | Degree_days_FM |  | °C day<sup>-1</sup> | FLO to FM |
|  | Degree_days_Cycle |  | °C day<sup>-1</sup> | Emergence to FM |
| Geographic variables | Latitude | Latitude coordinate | degrees | Trial specific |
|  | Longitude | Longitude coordinate | degrees | Trial specific |
|  | Altitude | Altitude (above sea level) | m | Trial specific |
| Management variable | emd | crop emergence date | Julian days | Trial specific |

The step of collections involves the use of different databases. The climate data set was collected from the weather station from INMET (Brazilian Institute of Meteorology - https://portal.inmet.gov.br/), located at the trial municipality. For trials with no weather stations available in the municipality, we used daily gridded climate data from NASA POWER (Sparks, 2018).

After relating trials to climate data, the step of data processing was conducted by adopting different sampling levels of environmental information and then translating them into actual environmental covariates (hereafter “EC”) capable of better capturing temporal variation across the crop life cycle. The development stages were computed at a field trial level using the mean values of FD and MF as observed in each trial. For the reproductive stage, according to the information supplied by rice breeders from Embrapa Rice & Beans, we assumed that panicle initiation (PI, corresponding to the stage R0 described by Counce et al., 2000) began 25 days before FD, as PI is not observed at field trials. The effective daily heat units (Degree days) were calculated based on daily mean temperature and three cardinal temperatures: base (8ºC), optimum (30ºC), and maximum (42ºC) thresholds. Details on the equation of degree days is described in Heinemann et al. (2017) and Bouman et al. (2001). We adopted this strategy because we focused on screening EC over the irrigated rice’s GY in the Brazilian tropical region. The sampling levels adopted in this study were:

A. Crop cycle: sampling of EC during the period between emergence date (EMD) and maturation date (MD), that is, involving maximum, minimum, mean, and accumulated values for each EC across the whole crop life cycle (V1 to R8);
B. Vegetative phase: sampling of maximum, minimum, mean, and accumulated values for each EC only for the vegetative period (from EMD to PI (V1 to R0));
C. Reproductive stage (R): sampling of maximum, minimum, mean, and accumulated values for each EC from PI to FD (R1 to R4);
D. Panicle initiation (PI): sampling values for EC only for PI date (R0 stage), defined here as 25 days before FD;
E. Flowering period (FLO): mean sampling values for EC only for the flowering period, i.e., four days before and four days after the flowering date (R4 stage), defined here as more than 50% of plants with open flowering; and
F. Filling grain stage (FG): sampling of maximum, minimum, mean, and accumulated values for each EC after FD until MD (R5 to R9 stages).

As the objective of this study is to provide a broader assessment of environmental covariables relating them to irrigated rice yields, considering vegetative, reproductive, and filling grain stages, in the Brazilian tropical irrigated rice production region (Midwest, North, Southeast, and Northeast megaregions), we averaged genotype yields per trial. Figure S1 (supplementary information) shows the coefficients of variation (CV) for trials across megaregions per genotype groups “IRGA 424,” “BRS Catiana,” and “All genotypes.” The CVs are below 30%, which is an evidence for the representativeness of trial average values used in this study (Figure S1, supplementary information). Finally, the final matrix of ECs comprises 38 climatic variables, three geographic factors, and one management variable. Table 1 shows all the detailed information.

### 2.5 Learning the empirical “environmental clusters” from yield data

Using historical yield data, we searched for independent environment clusters considering the groups “All genotypes,” “IRGA 424,” and “BRS Catiana.” We used a method of unsupervised learning aiming to perform tasks such as classification. Towards this aim, we applied “K-means,” an unsupervised machine learning (ML) algorithm for pattern recognition. The number of clusters (environments) was defined based on the following criteria: a) Elbow method (based on the within cluster sums of squares), and b) empirical expertise and knowledge of irrigated rice breeders about the target production region. The similarity between any two pieces of data was quantified in terms of Euclidean distance (Hartigan and Wong, 1979). The analysis was performed using the Sklearn.learn library (Pedregosa et al., 2011) in Python programming language, v. 3.7 (Van Rossum and Drake, 2009).

### 2.6 Envirotyping-based prediction of environmental clusters

After labelling the observed yield levels in empirical clusters (“environmental clusters”) using the “K-means” algorithm, we then applied a second supervised learning method to associate this “environmental clusters” with the actual environmental information (EC matrix). For this, we adopted a nonparametric unsupervised learning method used for classification and called it “decision tree models” (DTC), considering each genotype group as previously described (“IRGA 424,” “BRS Catiana,” and “All genotypes”).

DTC was chosen because it is an efficient classification tool that presents several advantages, such as (1) it is computationally inexpensive, (2) makes no assumptions about the distribution of ECs, and (3) is robust regarding redundant ECs (Tan et al., 2014; Beucher et al., 2019). The target variable was the observed yield clustered (environments) by “K-means,” and the features variables was the EC (independent or exploratory variables; see Table 1). The objective was to identify ECs that are discriminants for changing in environments (clusters) for each group (“IRGA 424,” “BRS Catiana,” and “All genotypes”) and its respective transition probabilities. The DTC estimation process takes into account the amount of information available in the database (the information needed to estimate a tree of depth d is approximately 2^d^). Based on that, for each genotype group (“IRGA 424,” “BRS Catiana,” and “All genotypes”), we applied the entire database looking for a model that better predicted the association between ECs and the clustered yield (environments). The EC selection that composes the final DTC was chosen by the DecisionTreeClassifier function from the Sklearn.learn library (Pedrosa et al., 2011), which selects ECs according to the index of “gini impurity.” To avoid overfitting, the tree control size (max_depth) was set at 5 (five levels) for all genotype group data sets (“IRGA 424,” “BRS Catiana,” and “All Genotypes”). Statistical measurements of performance are given by equations 1 and 2, such as accuracy (eq. 1), precision (eq. 2), and recall (eq. 3) (fraction of TP from the total amount of TP and false negatives (FN)):

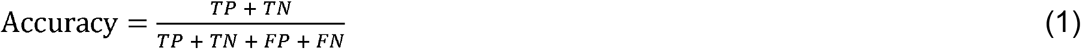

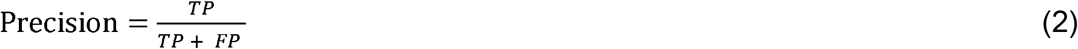

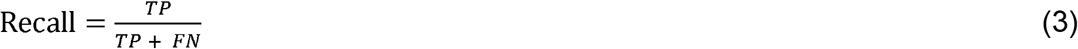

where TP means True Positive, which is equivalent to the number of cases correctly identified as positive; FP means False Positive, which is the number of cases incorrectly identified as positive; TN means True Negative, which is the number of cases correctly identified as negative; FN is False Negative, which is the number of cases incorrectly identified as negative. As there are unbalance classes (clusters), to calculate recall we weighted the averages among all classes. The analysis was performed using Python programming language, v. 3.7 (Van Rossum and Drake, 2009).

### 2.7 Typology discovery

In order to better dissect and understand DTC results, we created scenarios considering the median and the cut values of quantiles 1 (Q1, 25%), 2 (Q2, 50%), and 3 (Q3, 75%) of the most important EC variables discriminated by DTC for each group (“IRGA 424,” “BRS Catiana,” and “All genotypes”). The process of typology discovery (Costa-Neto et al., 2021) involves the expansion of these scenarios and the combination of the most important ECs and their values in the quantiles 25%, 50%, and 75%, used as thresholds for discretizing the continuous distribution of environmental factors and finding the frequency of the predominant “environmental types” (Clusters) for the most important ECs.

### 2.8 Effectiveness of environmental clusters in supporting GE analysis

The effectiveness of environmental clustering in supporting GE analysis was verified in two steps. First, two statistical models were fitted considering the grain yield of 34 advanced yield trials (AYT) conducted between 2011 and 2017. 76 genotypes were considered for this setting. Two liner mixed models (LMM) were fitted: the first considered the sources of variation due to location (L), year (Y), genotype (G), and interactions (GL, GY, LY, GLY) as follows:

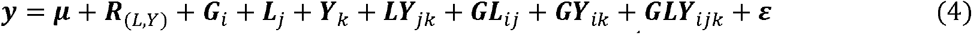

where: ***y*** is a vector of phenotypic observations (grain yield, kg ha^-1^) accounting for i-genotypes, j-locations, k-years; ***R***_(*L,Y*)_ is the fixed-effect vectors for replications (r = 1,…,4) within locations and years; ***G***_*i*_ is vector of random effects for *i*-genotypes, with 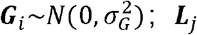 is a vector of random effects for *j*-locations, with 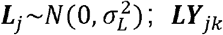 is vector of random effects for the interactions between *j*-locations and *k*-years, with 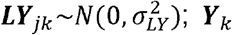 is vector of random effects for *k*-years, with 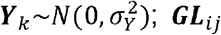 is vector of random effects for genotypes by location interactions, with 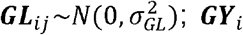 is vector of random effects for genotypes by year interactions, with 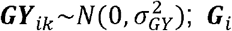 is vector of random effects for the interactions genotype x location x year, with 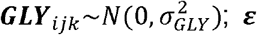 is vector of residual random effects, with 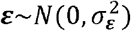.

From the LMM outcomes, the genotypic correlation across locations and years were computed as: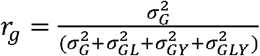, as a measurement of the broad genotypic correlation for environments (or ratio between broad G and GE effects), 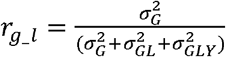, as the genotypic correlation across locations in any year,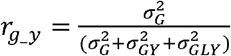 as the genotypic correlation across years at any location, 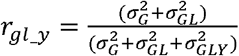 as the genotypic correlation across locations for a given year, and 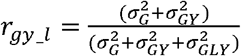 as the genotypic correlation across years for a given location. As a reference representing GE pattern of those correlations, we adopted the criteria obtained from simulation studies that suggests a predominant crossover GE pattern (complex) when *r* < 0.2, and a predominant non-crossover GE pattern (simple) when *r* > 0.8 (Cruz and Castoldi, 1991).

A second LMM was fitted accounting for clusters variation. From (4), we added the effect of yield clusters in two ways: (1) by splitting the genetic variation (G) into a broad genetic effect plus cluster-specific variation, and (2) by splitting environmental variations (locations, years) into within and across clusters. Consequently, the model (4) is now:

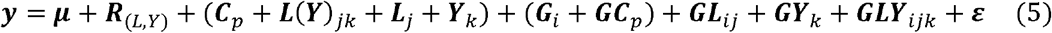

where *C*_*p*_ is the random cluster effect, with 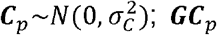 is the variation of locations, with 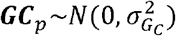. Thus, the genetic variance component is the sum of the effects (***G***_*i*_ +***GC***_*p*_), which consequently changes the denominator of all previous described correlation equations, now accounting to the component 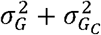.

Finally, we aimed to validate the effectiveness of clusters across a wide number of experimental network structures. For this, we ran year-specific LMM considering a random set of genotypes and locations sampled from the original data set. For this, we adopted a permutation scheme to simulate 5,000 scenarios of experimental networks for each year. To avoid possible noises due to sample size, we considered only the years with a higher number of environments (2011, 2012, 2013, and 2017). For each permutation, we considered a sample of 70% of genotypes and 90% of environments for a given year, resulting in 20,000 moment estimations of variance components related to genotypic and genotype by location interaction. As each LMM was fitted considering within-year variations, the models (4) and (5) were written as:

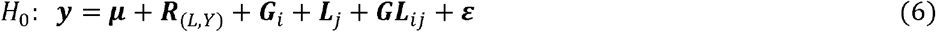

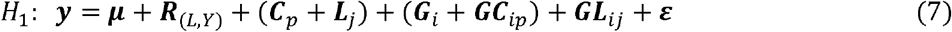

Where *H*_0_ is the reference model (null-hypothesis) and *H*_1_ is the updated model with the effect of clusters for controlling the environmental variability across locations (alternative hypothesis). The resulted distribution of genetic correlations (n = 20,000) for each model was compared using the Kruskal-Wallis nonparametric method. Consequently, the estimation of year-related correlations was not computed; only the location and GL within each year were considered. The LMM approach was implemented in R using the package *lme4* (Bates et al., 2015).

## 3 RESULTS

### 3.1 Regional variations in grain yield in Brazil

The average yield of tropical irrigated rice in Brazil is 6,224 kg ha^-1^ considering all genotypes in all study years (Figure 2A). However, the distribution of yield of irrigated rice across each tropical production region in Brazil highly depends on geographic specific conditions (Figure 2). In spite of this, possible seasonal effects led to an increase in diversity of yield levels in some States. The average yield value for the Midwest (MW, red color, 5,539 kg ha^-1^) and North (NO, green color, 5,492 kg ha^-1^) is almost the same, while yield distribution for different States ranges in magnitude. For the Goiás State (GO), where the nursery of the EMBRAPA breeding program is, located, the highest diversity of yield levels is expected, surpassing almost all yield levels found in the other regions. This also occurs for the Tocantins State (TO) in the NO region, in which the higher instability of production characterizes this State as the less predictable and most prone to yield losses. However, this may be associated with the higher number of locations considered for these two states in this study (TO = 52 field trials, GO = 47 field trials). The Northeast (NO) seems to be the region where there are the highest yield values, except for the States of Sergipe (SE) and Alagoas (AL). Higher productions were also observed in São Paulo State (SP, Southeast), but the resolution of this yield level might be low due to the small sample size (SP = 3 field trials).

**Figure 2.**
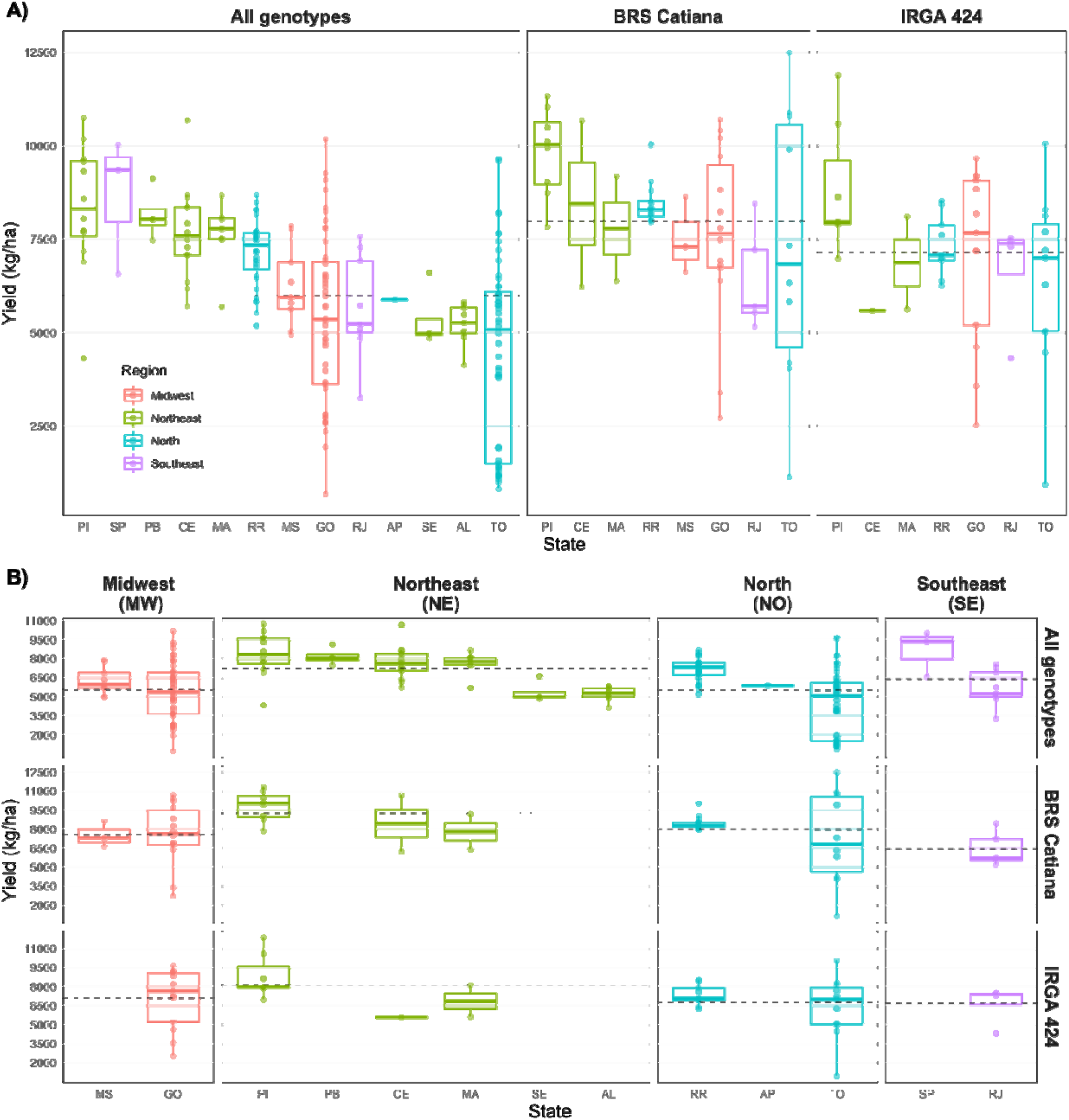
Empirical characterization of the yield levels in tropical irrigated rice for each region (Midwest, Northeast, North and Southeast) and genotype group (All genotypes, BRS Catiana and IRGA 424) in Brazil. **(A)** Boxplots of grain yield (kg/ha) for each genotype group at each state. **(B)** Boxplots of grain yield divided by each region and genotype group. Horizontal dotted lines denotes the average yield value for the respective panel.

In a same distribution of tropical conditions, the BRS Catiana outperforms IRGA 424 and the other genotypes (Figure 2). The average yield level across region is 7,980 kg ha^-1^, ranging from 6,416 kg ha^-1^ in the SE region to 9,246 kg ha^-1^ in the NE region, whose production is almost the same in the regions MW (7,581 kg ha^-1^) and NO (7,955 kg ha^-1^). The cultivar IRGA 424 also outperforms the other genotypes, with an average yield production in Brazil of 7,153 kg ha^-1^, ranging from 6,657 kg ha^-1^ and 6,812 in the SE and NO regions, respectively, to 7,085 kg ha^-1^ and 8,120 kg ha^-1^ in the NO and MW regions, respectively. Thus, it seems that there is no cultivar x region interaction for grain yield production, which supported our approach of running a joint nationwide model accounting for historical and climatic data across all tropical region.

### 3.2 Pattern-recognition of environmental cluster quality scenarios

As the empirical observation of yield distribution does not provide easily interpretable results, we used a K-means algorithm to pinpoint clusters of yield levels (environments) across regions. This analysis was conducted considering all genotypes (“All genotypes” group) and cultivar-specific groups (“BRS IRGA” and “BRS Catiana”). Supplementary Figure S2 shows the Elbow method (based on the within cluster sums of squares), one of the criteria used to select the cluster number for each genotype group. It allowed us to identify four environmental clusters (yield clusters). They were denominated after (environment 1) Very low yield, (environment 2) Low yield, (environment 3) High yield, and (environment 4) Very high yield (Table 2, Figure 3).

**Table 2.** Summary of the grain yield distribution of the four yield clusters founded for irrigated rice in Brazil, considering each genotype group.

| Group | Cluster | Statistic |  |  |  |  |
| --- | --- | --- | --- | --- | --- | --- |
|  |  | Average<br>(kg.ha<sup>-1</sup>) | Range<br>(kg.ha<sup>-1</sup>) | Deviation<br>(kg.ha<sup>-1</sup>) | CV<br>(%) | # of Location<br>(Frequency, %) |
| **All**<br>**genotypes** | (1) Very Low Yield | 1773 | 688 - 3255 | 732 | 41 | 28 (14 %) |
|  | (2) Low Yield | 5104 | 3513 - 6088 | 733 | 14 | 71 (35 %) |
|  | (3) High Yield | 7192 | 6163 - 8085 | 566 | 8 | 74 (36 %) |
|  | (4) Very High Yield | 9047 | 8168 - 10751 | 773 | 9 | 30 (15 %) |
| **BRS Catiana** | (1) Very Low Yield | 3102 | 1146 - 4200 | 1241 | 40 | 5 (9 %) |
|  | (2) Low Yield | 6432 | 5158 - 7330 | 658 | 10 | 15 (26 %) |
|  | (3) High Yield | 8372 | 7491 - 9181 | 446 | 5 | 22 (39 %) |
|  | (4) Very High Yield | 10581 | 9713 - 12493 | 702 | 7 | 15 (26 %) |
| **IRGA 424** | (1) Very Low Yield | 2347 | 940 - 3573 | 1325 | 56 | 3 (6 %) |
|  | (2) Low Yield | 5346 | 4324 - 6375 | 740 | 14 | 11 (23 %) |
|  | (3) High Yield | 7674 | 6938 - 8633 | 529 | 7 | 25 (53 %) |
|  | (4) Very High Yield | 9812 | 8841 - 11891 | 1018 | 10 | 8 (17 %) |

**Figure 3.**
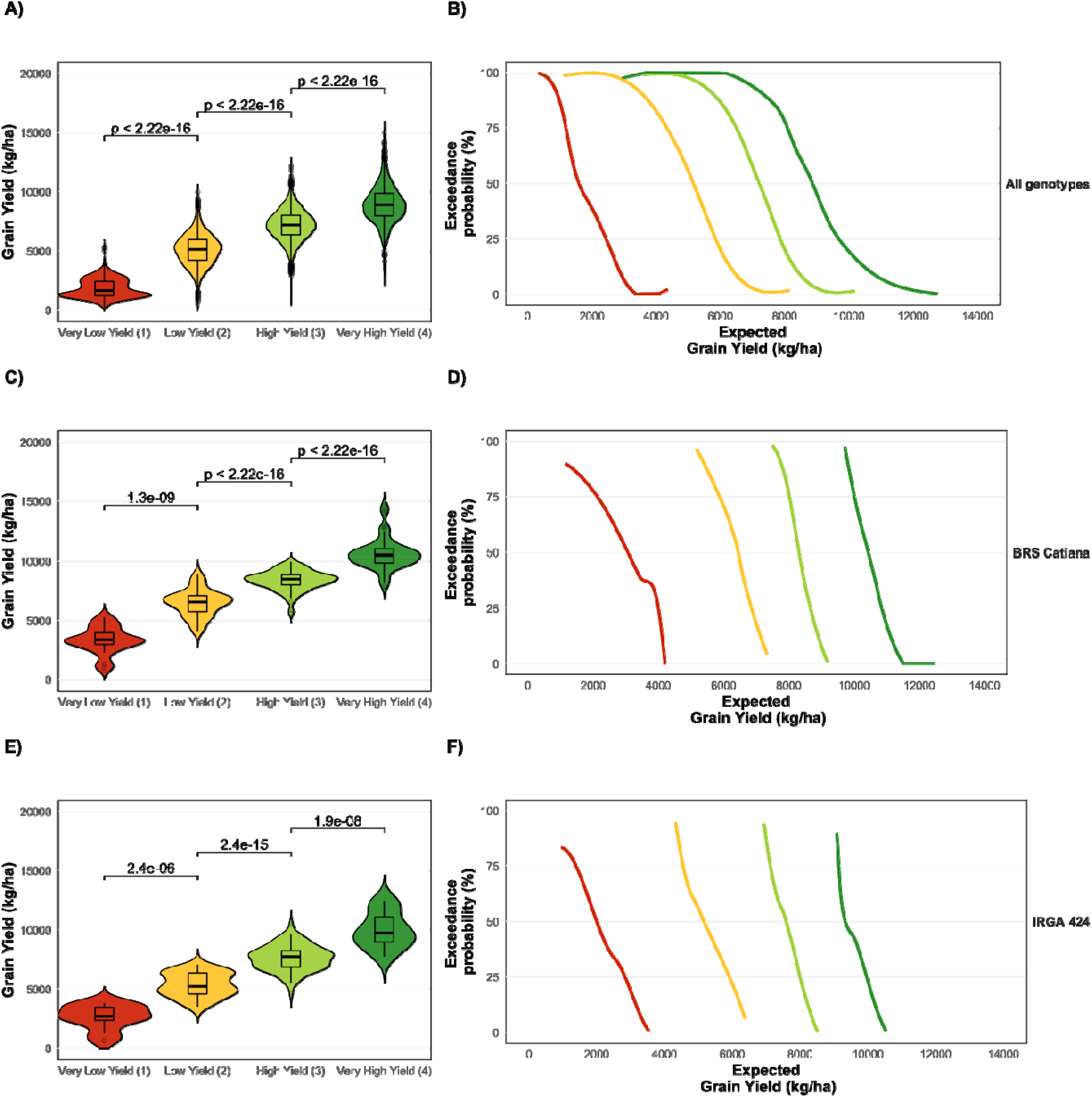
Discriminant power of the pattern-recognition approach in identifying clusters of environmental quality for grain yield in irrigated rice. (**A**) violin-plot considering the group of all genotypes (“All genotypes”) for each yield cluster. (**B**) Exceedance probability for the group of all genotypes; (**C**) violin-plot considering the cultivar-specific group of “BRS Catiana” or each yield cluster. (**D**) Exceedance probability for the cultivar-specific group of BRS Catiana; (**E**) violin-plot considering the cultivar-specific group of “IRGA 424” for each yield cluster. (**F**) Exceedance probability for the cultivar-specific group of IRGA 424. p-values denotes the significance from the Kruskall-Wallis test.

Figure 3 shows the discriminant power of the approach. There were significant differences among clusters (environments) for all genotype groups (from Kruskall-Wallis method, p < 0.001). For the cultivar-specific groups (BRS Catiana and IRGA 424), the divergence among groups is even greater than those observed considering all genotypes in the irrigated rice germplasm.

The lower yield cluster (environment 1) is the more unstable and less productive cluster, but it is also the less frequent for all genotype groups, such as “all genotypes” (n = 5 locations, 9% of the targeting region), BRS Catiana (n = 28 locations, 14% of the targeting region for this cultivar), and IRGA 424 (n = 3 locations, 6% of the targeting region for this cultivar). The clusters (environment 2) and (environment 3) are the most frequent in all groups of this study. Together, they cover more than 71% of the targeting area for the group “All genotype,” 37% for BRS Catiana, and 36% for IRGA 424. Finally, the coefficient of variation (CV, %) decreases as the yield level increases, which suggests that the clusters (environment 3) and (environment 4) are also the most productive and stable across years.

Figure 4 shows the geographic distribution and probability of occurrence of the four environmental clusters ((1) Very low yield; (2) Low yield; (3) High yield and; (4) Very high yield) and Figure 5 shows their regional frequency.

**Figure 4.**
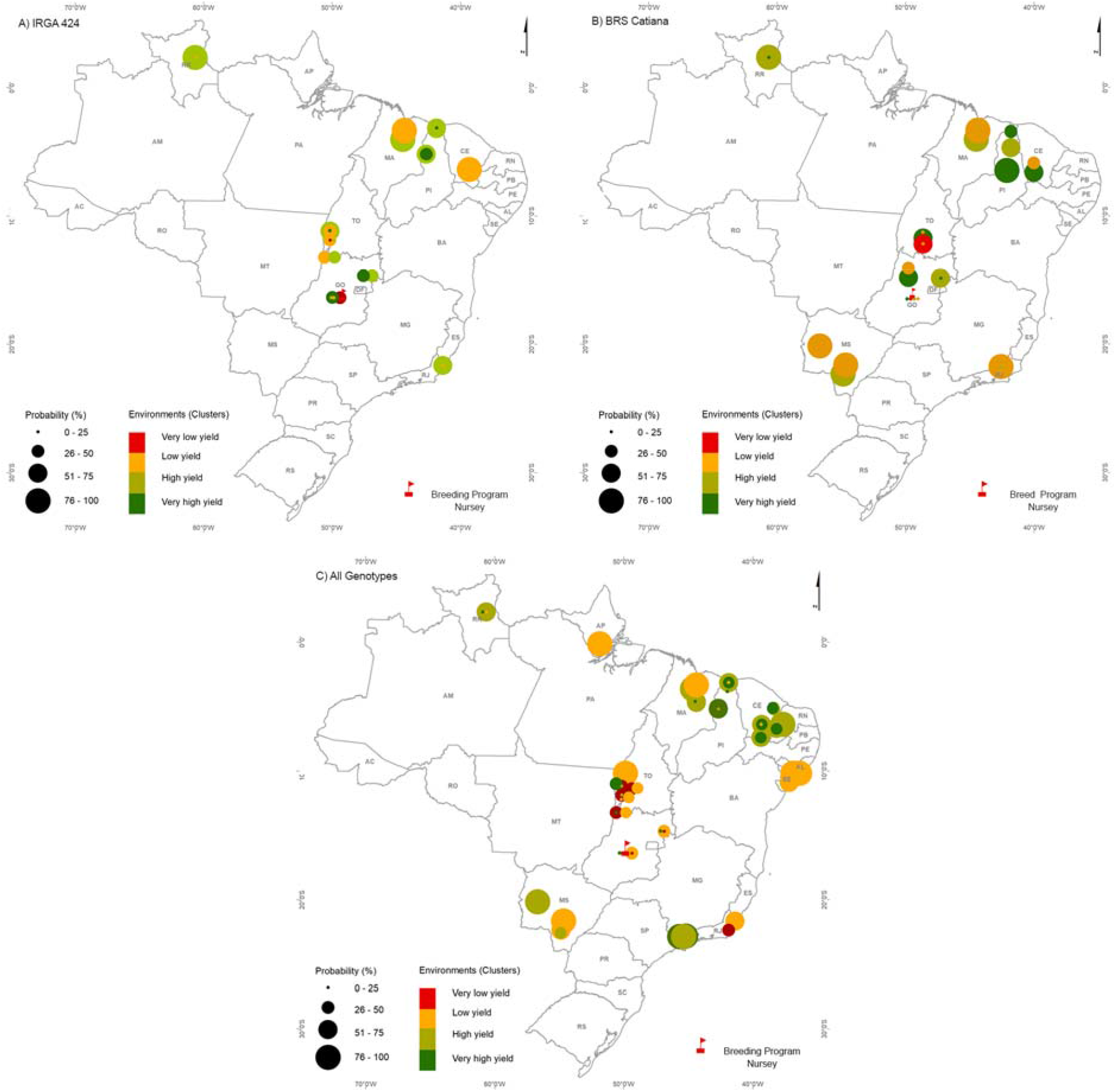
Geographic location of environments (Very low yield (1), red; Low yield (2), orange; High yield (3), light green and Very high yield (4), dark green) and their respective probability of occurrence (circle) across the tropical irrigated rice production region for genotypes groups. **(A)** IRGA 424; **(B)** BRS Catiana and **(C)** All genotypes. The circle diameter represents the probability of occurrence. The statistics of environments (Very low yield (1), red; Low yield (2), orange; High yield (3), light green and Very high yield (4), dark green) is described at Table 2.

**Figure 5.**
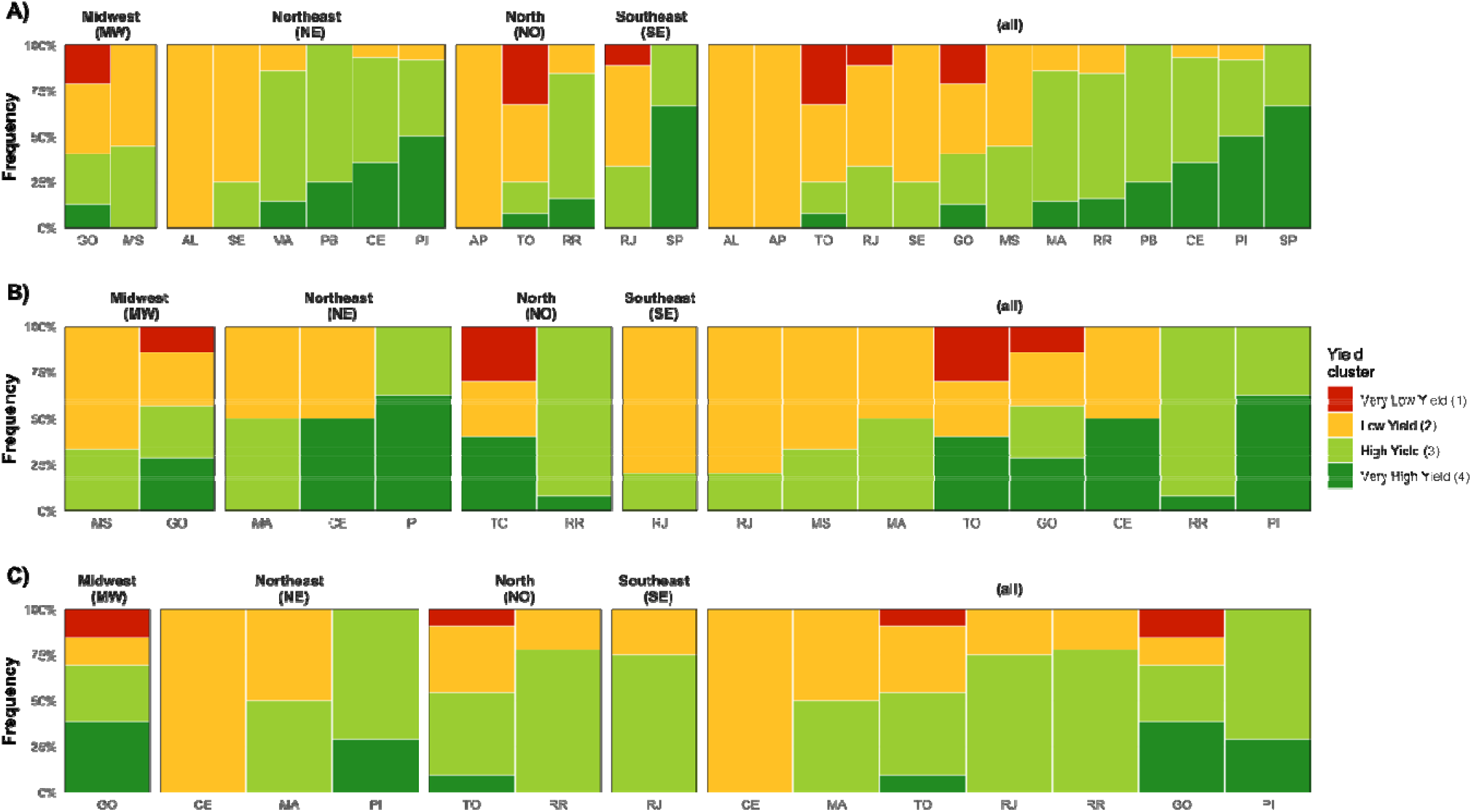
Frequency of each yield cluster across each Brazilian region and states. **(A)** panel of frequencies considering “All genotypes” group (n=471 genotypes and 203 trials); **(B)** panel of frequencies considering the cultivar-specific group “BRS Catiana” (n=1 genotype and 57 trials); **(C)** panel of frequencies considering the cultivar-specific group “IRGA 424” (n=1 genotype and 47 trials). Panels with “(all)” denotes the combination of all regions for a given genotype group.

### 3.3 Characterization of environmental cluster diversity

As the geopolitical discrimination of the regions seems to reflect some macro-variations in yield values, which may be associated to the geographic diversity of edaphoclimatic and biotic stress conditions, it is also possible to argue that most of the diversity of such differences could be due to seasonal (climatic) variations (Figure 4 and 5). Then, to study these seasonal variations, we decided to characterize the environments using additional descriptors of environmental diversity derived from the envirotyping protocol. In short, these descriptors are used as environmental covariables (EC), here scaled and centered (Z scores) to support a better visualization of similarity in further analyses (Figure 6A) and with its distributions across regions in actual scales (Figure 6B).

**Figure 6.**
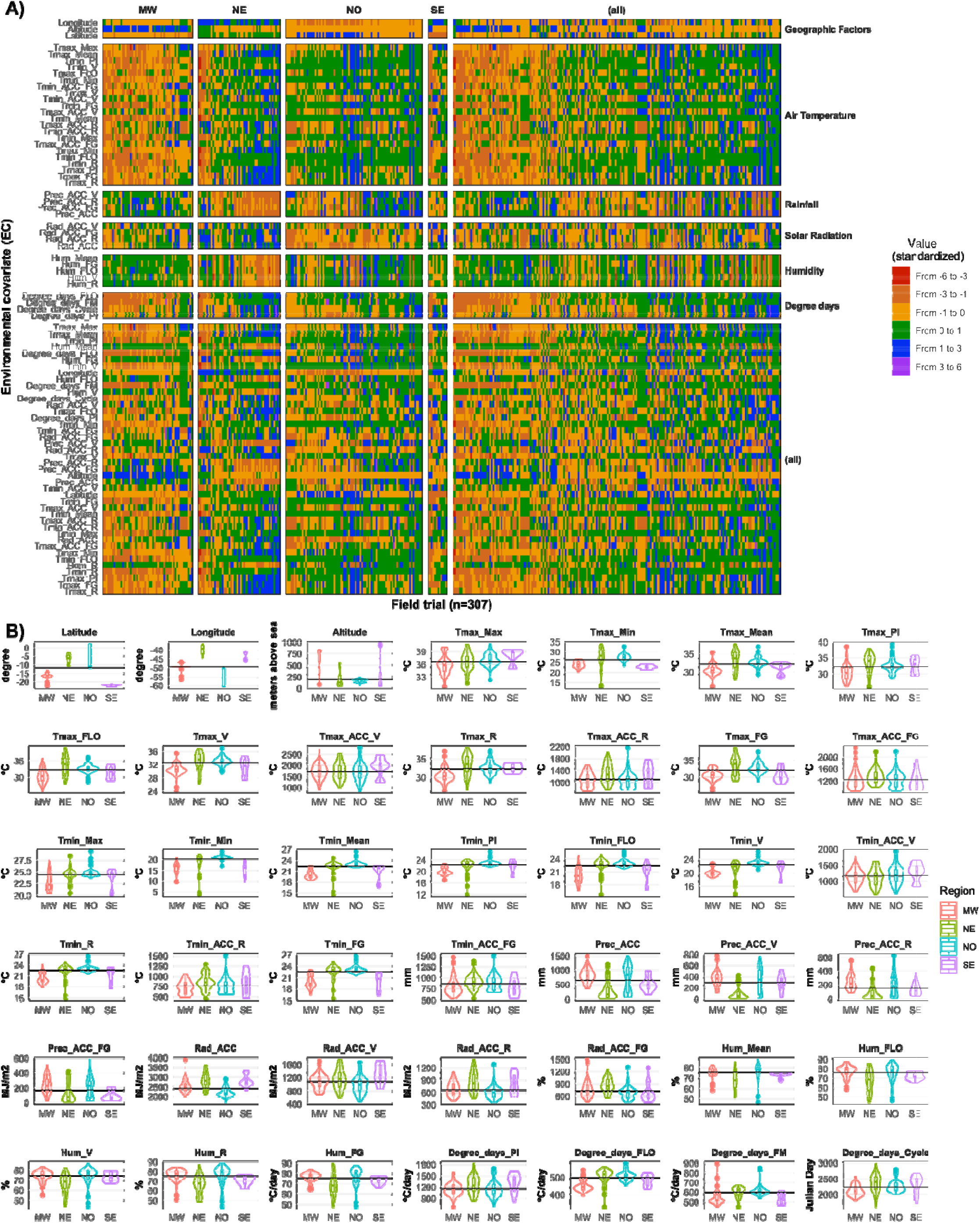
Complete macro-environmental characterization of the four production regions in Brazil (MW, Midwest; NE, Northeast; NO, North and SE, Southeast). **(A)** Panel of the main climatic and geographic covariates standardized (Z-score) and across each field trial (n=307) and region. **(B)** Distribution of each environmental covariable across regions represented by violin plots. Horizontal lines denotes the average value for each EC across all regions and years. Tmax_FLO. maximum temperature for flowering period; Tmax_Max. maximum value of maximum temperature; Tmax_Mean. mean value of maximum temperature; Tmax_Min. minimum value of maximum temperature; Tmin_FLO. minimum temperature in the flowering period; Tmin_Max. minimum value of maximum temperature; Tmin_Mean. mean value of minimum temperature; Tmin_Min. minimum value of minimum temperature; Degree_days_Cycle. degree days accumulated; Prec_ACC. precipitation accumulated; Radiation_ACC. solar radiation accumulated. Tmax_Acc. maximum temperature accumulated and Tmin_ACC. minimum temperature accumulated.

The Northeast (NE) is characterized by higher values of temperature, especially maximum temperatures during the crop cycle and the flowering period (Figure 6). Due to its geographic position, this region is also characterized by higher longitude values and lower latitude values. There is a lower diversity of elevation levels (altitude), which differs from other regions such as Southeast (SE) and the Midwest (MI), where there is the highest diversity of elevation. Finally, NE is also characterized by higher values of solar radiation in most of the locations of this study. NE has also the lowest accumulated rainfall among all regions, especially during vegetative stages (V). At this region, the most commonly found yield level is very high yield (dark-green) and high yield (green) for “all genotypes” and the group “BRS Catiana,” except for “IRGA 424,” which seems not to be adapted for this region (Figure 4 and 5).

The North (NO) region has a common pattern of low elevations and global solar radiation (Figure 6). In almost every field trial, the values of growing degree days of almost every development stage were higher than those of any other region, especially long reproductive and filling grain phases, despite having almost the same cycle duration than other places do. This region has two different rainfall regimes: one higher in the States of AP and RR, and lower rainfalls in the TO State. Despite this, the patterns of humidity and temperature are diverse within this region. Considering every genotype group studied in this work, it seems that NO is very diverse in terms of yield levels, where the TO State is very diverse inside itself (Figure 4 and 5). The Roraima State (RR) seems to be the most stable regarding production and presenting the highest yield levels, while Tocantins has regions more prone to loess and lower yield levels. Despite this, the TO state also has a higher probability of achieving higher yield levels, which suggests the importance of seasonal factors in defining yield levels.

While the NO region has the highest values of accumulated rainfall and the lowest growing degree days, the Southeast (SE) region showed the highest variation for degree days and the poorest rainfall regimes. This is one of the most diverse regions, despite being the region with the fewest observations. The yield levels observed in SE (Figure 4 and 5) show that most of this diversity comes from the Rio de Janeiro State (RJ), with a predominance of low yield levels (yellow color in Figures 4 and 5), while the São Paulo State (SP) has a predominance of higher yield levels. For the groups BRS Catiana and IRGA 424, it seems there is a State x cultivar interaction, where low yield levels are more frequent for BRS Catiana while higher yields are more frequent for IRGA 424.

The Midwest (MI) is characterized by patterns of lower temperatures and increasing degree days while there is a predominance of higher elevations and air humidity (Figure 6). For irrigated rice, it seems to be very stable regarding climatic factors. However, this State (Figure 4 and 5) has the highest diversity of yield levels, in which every yield cluster occurs in almost the same frequency as in Goiás State (GO), where the breeding program nursery is. This is a repeatable pattern in every genotype group studied.

### 3.4 Decision tree model (DTC) performance and scenarios

Yield cluster levels associated with additional information derived from environmental covariables (ECs). The decision tree classification models (DTC) showed an accuracy (precision and recall) of 93.6 (94.9 and 93.6), 91.2 (93.2 and 91.2), and 82.3 (84.3 and 82.2%) for the genotype groups “IRGA 424,” “BRS Catiana,” and for “All genotypes,” respectively. The DTC precision for yield clusters (homogeneous environments) in each genotype groups (“IRGA 424,” “BRS Catiana,” and “All genotypes”) are illustrated in the confusion matrix (Supplementary Figure S3).

For a better understanding of DTC results (Figures 7 and 8), we constructed scenarios considering the median and the cut values of quantiles 1 (Q1), 2 (Q2), and 3 (Q3) for each genotype group described in Table 3. For median and Q1, Q2, and Q3 scenarios, ECs (environmental covariables) are fixed at median, Q1 (25%), Q2 (50%), and Q3 (75%) values, respectively.

**Figure 7.**
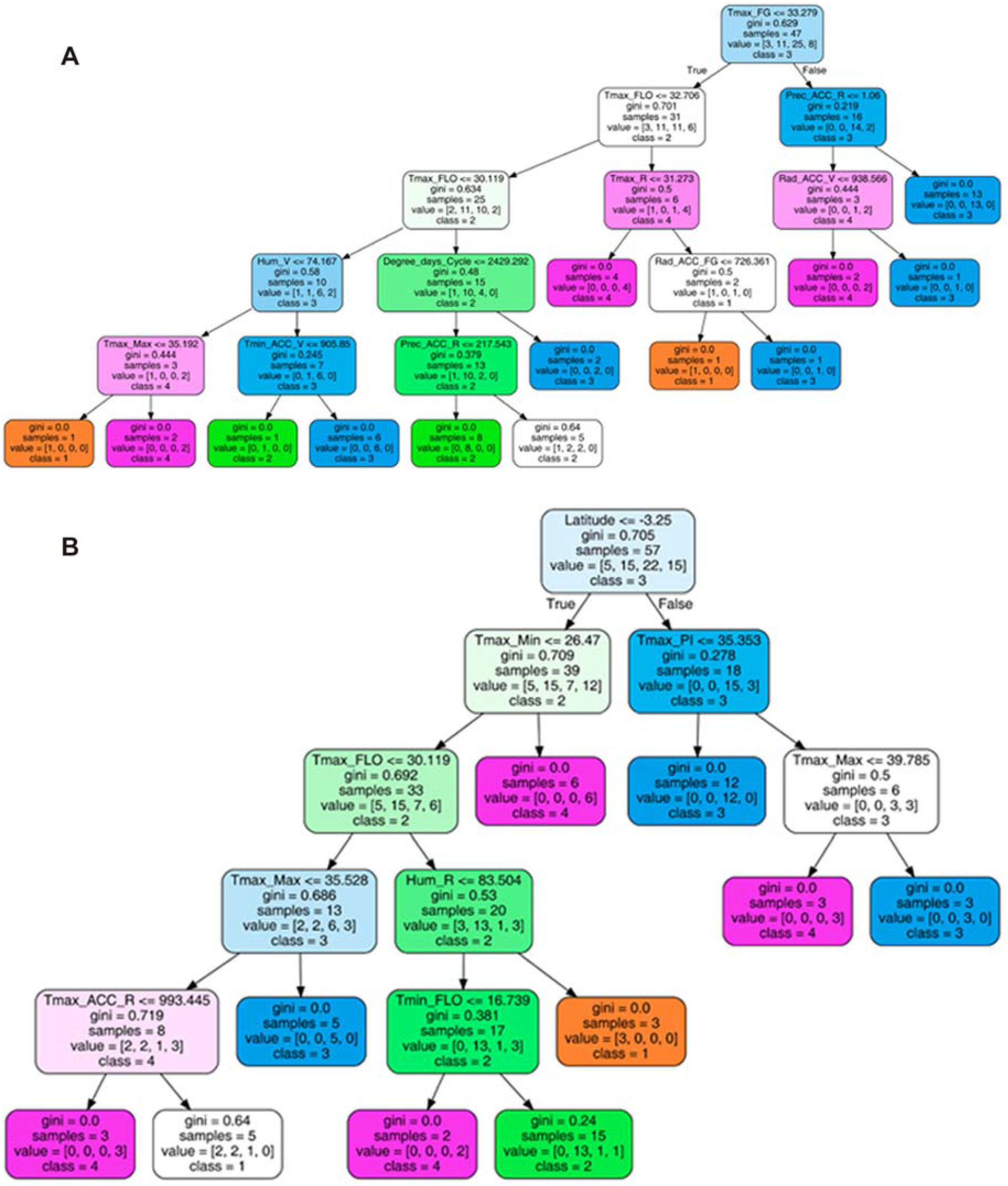
Decision tree for the analysis of environment covariables of influence on the irrigated rice yield for genotype groups. **(A)** “IRGA 424” and **(B)** “BRS Catiana”. In each rectangle, in the top row, the most significant environmental covariable (EC) selected and their cut-off point (the value of the variable in which the division is made) are given; the second line represents the gini value; the third line the number of sample, the fourth line the value (sample distribution in the environments [Very low yield (1); Low yield (2); High yield (3) and Very high yield (4)] and class represents the environments (1) Very low yield; (2) Low yield; (3) High yield and (4) Very high yield].

**Figure 8.**
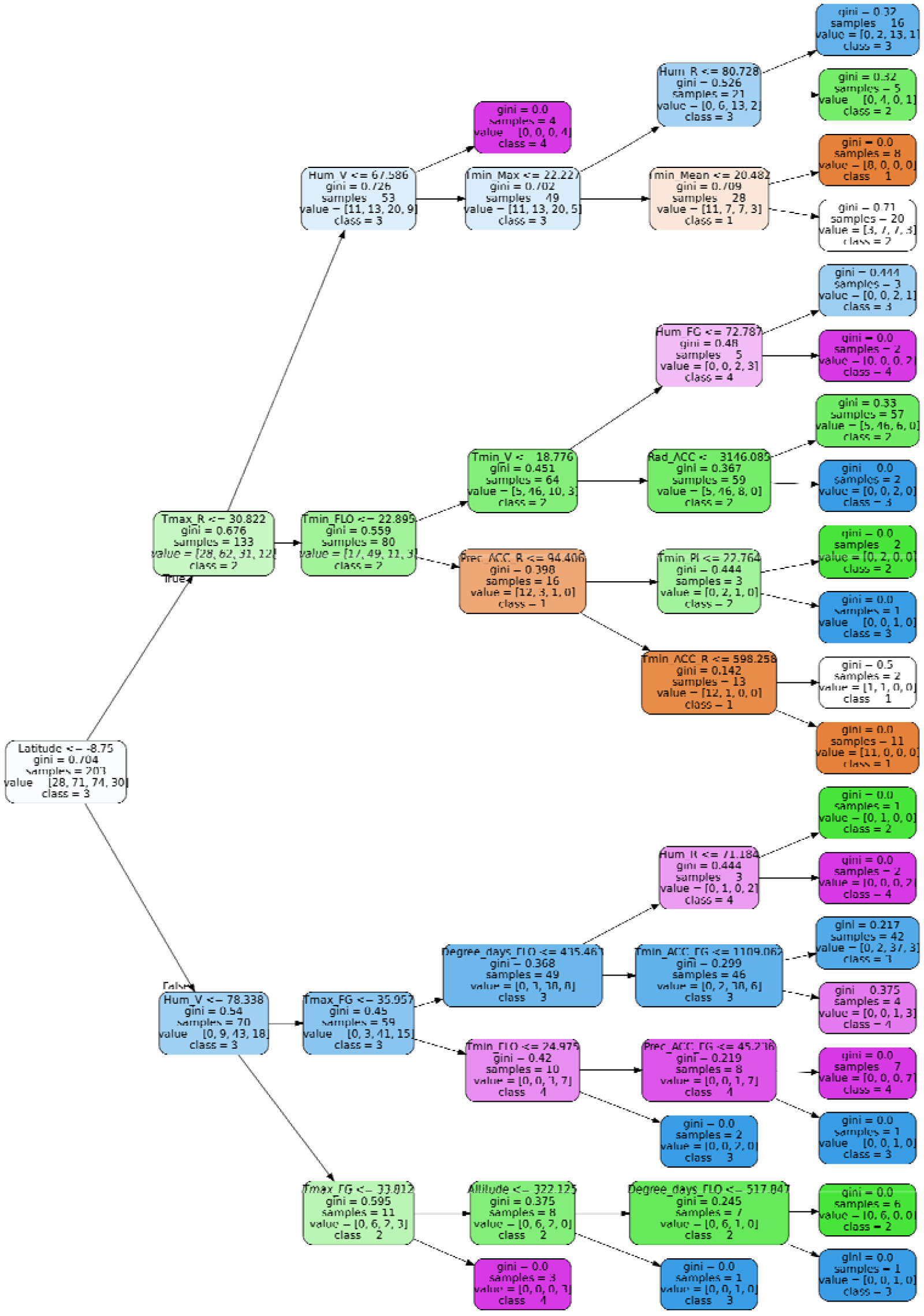
Decision tree for the analysis of environment covariables of influence on the irrigated rice yield for genotype group “All genotypes”. In each rectangle, in the top row, the most significant environmental covariable (EC) selected and their cut-off point (the value of the variable in which the division is made) are given; the second line represents the gini value; the third line the number of sample, the fourth line the value (sample distribution in the environments [Very low yield (1); Low yield (2); High yield (3) and Very high yield (4)] and class represents the environments (1) Very low yield; (2) Low yield; (3) High yield and (4) Very high yield].

**Table 3.** Scenarios considering the values of environmental covariables discriminated by the tree decision model (DTC, Figure 7 and 8) fixed at median and quantiles 1 (Q1, 25%), 2 (Q2, 50%) and 3 (Q3, 75%) for genotyping groups “IRGA 424” (Figure 7A), “BRS Catiana” (Figure 7B) and “All genotypes” (Figure 8).

| Environmental<br>Covariable (Unit) | Values for “IRGA 424” |  |  |  |
| --- | --- | --- | --- | --- |
|  | median | Q1(25%) | Q2(50%) | Q3(75%) |
| Tmax_FG (°C) | 32 | 30 | 31 | 35 |
| Tmax_FLO (°C) | 33 | 30 | 32 | 34 |
| Hum_V (%) | 70,1 | 66,3 | 71,7 | 77,1 |
| Tmax_Max (°C) | 37,3 | 35,8 | 36,9 | 39,1 |
| Tmin_ACC_V (°C) | 1160 | 1013 | 1158 | 1321 |
| Degree_days_Cycle (°C day<sup>-1</sup>) | 2171 | 2074 | 2170 | 2247 |
| Prec_ACC_R (mm) | 132 | 17 | 118 | 199 |
| Tmax_R (°C) | 32 | 30 | 31 | 34 |
| Rad_ACC_FG (MJ m<sup>-2</sup>) | 737 | 613 | 726 | 870 |
| Rad_ACC_V (MJ m<sup>-2</sup>) | 1026 | 837 | 970 | 1183 |
|  | Values for “BRS Catiana” |  |  |  |
|  | median | Q1(25%) | Q2(50%) | Q3(75%) |
| Latitude (degrees) | -9.18 | -16.35 | -11.15 | -3.15 |
| Tmax_Min (°C) | 26 | 24 | 26 | 28 |
| Tmax_FLO (°C) | 32 | 30 | 33 | 34 |
| Tmax_Max (°C) | 37 | 36 | 37 | 39 |
| Tmax_ACC_R (°C) | 1120 | 928 | 1104 | 1269 |
| Hum_R (%) | 70 | 64 | 73 | 80 |
| Tmin_FLO (°C) | 22 | 21 | 23 | 24 |
| Tmax_PI (°C) | 32 | 30 | 32 | 34 |
|  | Values for “All Genotypes” |  |  |  |
|  | median | Q1(25%) | Q2(50%) | Q3(75%) |
| Latitude (degrees) | -10.11 | -16.35 | -11.85 | -5.15 |
| Tmax_R (°C) | 32 | 31 | 32 | 33 |
| Hum_V (%) | 72 | 68 | 74 | 78 |
| Tmin_Max (°C) | 24 | 23 | 24 | 25 |
| Hum_R (%) | 72 | 67 | 75 | 80 |
| Tmin_Mean (°C) | 22 | 21 | 22 | 23 |
| Tmin_FLO (°C) | 22 | 21 | 23 | 23 |
| Tmin_V (°C) | 22 | 20 | 23 | 23 |
| Hum_FG (%) | 73 | 69 | 76 | 81 |
| Prec_ACC_R (mm) | 171 | 40 | 151 | 267 |
| Tmin_PI (°C) | 22 | 21 | 22 | 23 |
| Tmin_ACC_R (°C) | 781 | 607 | 772 | 890 |
| Tmax_FG (°C) | 32 | 30 | 32 | 34 |
| Degree_days_FLO (°C day<sup>-1</sup>) | 489 | 455 | 497 | 518 |
| Tmin_ACC_FG (°C) | 865 | 710 | 864 | 989 |
| Prec_ACC_FG (°C) | 183 | 64 | 167 | 282 |
| Altitude (m) | 316 | 126 | 197 | 519 |
| Rad_ACC (MJ m<sup>-2</sup>) | 2483 | 2141 | 2415 | 2722 |

For IRGA 424 (Figure 7A, Table 3), considering EC values of median (scenario median), the value of Tmax_FG (32 ºC, Table 3) are lower than the estimated value (33.279 ºC, Figure 7A); we then evaluated the next EC, Tmax_FLO. If we make a decision at this node, the DTC model predict two environmental clusters or classes, 2 (Low yield) and 3 (High yield), with the same probability (35.4%, Figure 7A). As the observed Tmax_FLO median value (33ºC, Table 3) is higher than the estimate value (32.706 ºC, Figure 7A), we must evaluate T_max_R. If we make a decision at this node, the DTC model predicts the environment 4 (Very high yield) with a probability of 67%. As the observed median value of Rad_ACC_FG (737 MJ, Table 3) is higher than the estimated value (726.361 MJ), the DTC model predicts the environment 4 (Very high yield) with 100% of probability. For the scenario Q1 (25%), considering EC values on Q1 (scenario Q1 - ECs fixed at 25%) and running the tree model until the last node, the DTC model predicts the environment 4 (Very high yield). For the scenario Q2 (50%, mean) and running the tree model until the last node, the DTC model predicts the environment 2 (Low yield). For the scenario Q3 (75%) and running the tree model until the last node, the DTC model predicts the environment 3 (High yield).

For BRS Catiana (Figure 7B, Table 3), considering EC values fixed on the median, the value of observed Latitude (−9.18) is lower than the estimated value (−3.25, Figure 7B). We then evaluated the next EC, Tmax_Min. If we make a decision at this node, the DTC model predicts the environment or class 2 (Low yield) (Figure 7B) with a probability of 38.4%. As the observed median value of Tmax_FLOR (32 ºC, Table 3) is higher than the estimated value (30.119 ºC), we evaluated the EC Hum_R. The observed value of Hum_R (74%, Table 3) is lower than the estimated value (83.504%, Figure 7B). We then evaluated Tmin_FLOR. The observed Tmin_FLOR value (22 ºC. Table 3) is higher than the estimated value (16.739 ºC, Figure 7B), therefore the DTC model predicts the environment 2 (Low yield) with a probability of 86.6%. For the scenario Q1 (25%) and running the tree model until the last node, the DTC model predicts the environment 1 (Very low yield). For the scenario Q2 (50%, mean) and running the tree model until the last node, the DTC model predicts the environment 4 (Very high yield). For the scenario Q3 (75%) and running the tree model until the last node, the DTC model predicts two environments, 4 (Very high yield) and 3 (High yield), with the same probability (100%).

For All genotypes (Figure 8, Table 3), considering EC values on the median, the value of observed Latitude (−10.11) is lower than the estimated value (−8.75, Figure 8). We evaluated the next EC, Tmax_R. If we make a decision at this node, the DTC model predicts the environment or class 2 (Low yield) with the same probability (46.6%). As the observed Tmax_R median value (32ºC, Table 3) is higher than the estimate value (32.822 ºC, Figure 8), we evaluated Hum_V. As the observed value of Hum_V (72%, Table 3) is higher than the estimated value (67.586%, Figure 8), the DTC model predicts the environment 4 (Very high yield) with a probability of 100%. For the scenario Q1 (25%) and running the tree model until the last node, the DTC model predicts the environment 4 (Very high yield). For the scenario Q2 (50%, mean) and running the tree model until the last node, the DTC model predicts the environment 4 (Very high yield). For the scenario Q3 (75%) and running the tree model until the last node, the DTC model predicts two environments, 3 (High yield) and 2 (Low yield).

### 3.5 Pinpoint environmental typologies across yield clusters

Figure 7 ((A) “IRGA 424” and (B) “BRS Catiana”) and Figure 8 (“All genotypes”) show the ECs discerned by DTC for each genotype group graphically. For the genotype group “IRGA 424,” the most discriminant ECs were maximum temperature at filling grain (Tmax_FG), maximum temperature at flowering period (Tmax_FLO), and accumulated rainfall at the reproductive phase (Prec_ACC_R). For the other levels of DTC, we observe that impurity (impurity - gini index) decreases, making the decision process more assertive. The lower the gini index, the lower the impurity associated to the classification power in a given node.

Figure 9 shows the frequency of these ECs (Prec_ACC_R, Tmax_FG and Tmax_FLO) across the environments (Very low yield (1), Low yield (2), High yield (3), and Very high yield (4)) for “IRGA 424.” It also shows the typology panels of the main environmental covariables. For the Very high yield (4) environment, a maximum temperature ranging from 31 to 32 ºC and 32 to 34 ºC at filling grain and flowering periods is expected in 50% of the occurrences, respectively (Figure 9B and C). Lower maximum temperatures at filling grain phase are expected at Very low (1) and Low (2) yield environments. Accumulated rainfall at the reproductive phase (Prec_ACC_R) ranged from 150 to 200 mm in 50% of the occurrences at Very High yield (4) environments (Figure 9A). Higher accumulated rainfalls (300 to 400 mm) are expected at Very low yield (1) environment in 50% of the occurrences.

**Figure 9.**
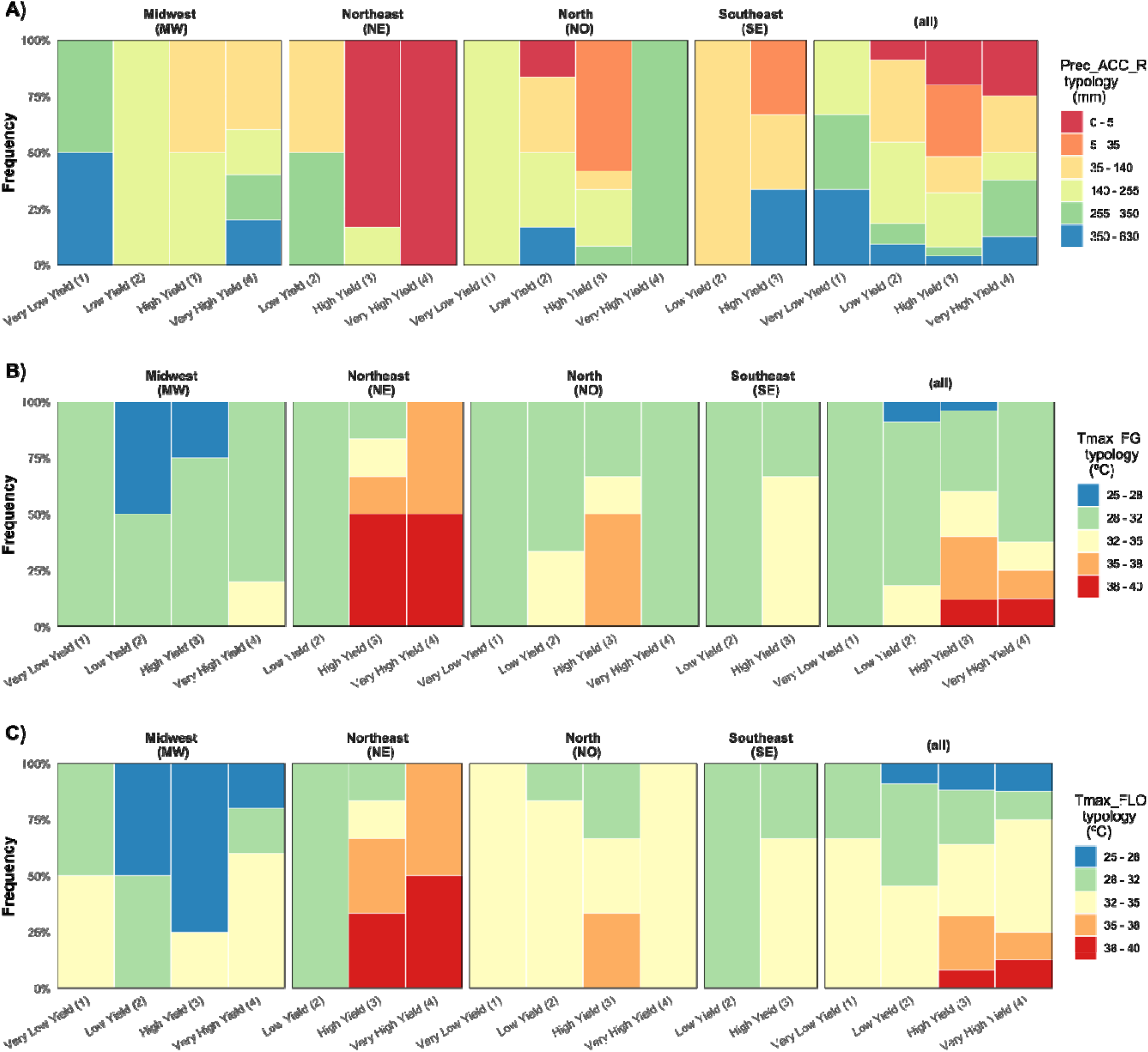
Global view of the frequency of occurrence of the main discriminants environmental covariate for genotype group “BRS IRGA” across the classifieds environments (clusters) **(A)** precipitation accumulated at reproductive phase (Prec_Acc_R) and **(B)** maximum temperature at filling grain (Tmax_FG) and maximum temperature at flowering (Tmax_FLO).

For “BRS Catiana,” the most discriminant ECs were Latitude, minimum value of maximum temperature for whole cycle (Tmax_min), and maximum temperature value for panicle initiation (Tmax_PI). Figure 10 shows the frequency of Tmax_min and Tmax_PI across the environments (Very low yield (1), Low yield (2), High yield (3), and Very high yield (4)). The minimum value of maximum temperature and the maximum temperature value at panicle initiation ranged from 24 to 26 and 30 to 32 ºC in 50% of the occurrences at Very high (4) yield environment, respectively. The worst environment, Very low yield (1), is characterized as having higher temperatures values for both Tmax_min and Tmax_PI considering 50% of the occurrences (Figure 10).

**Figure 10.**
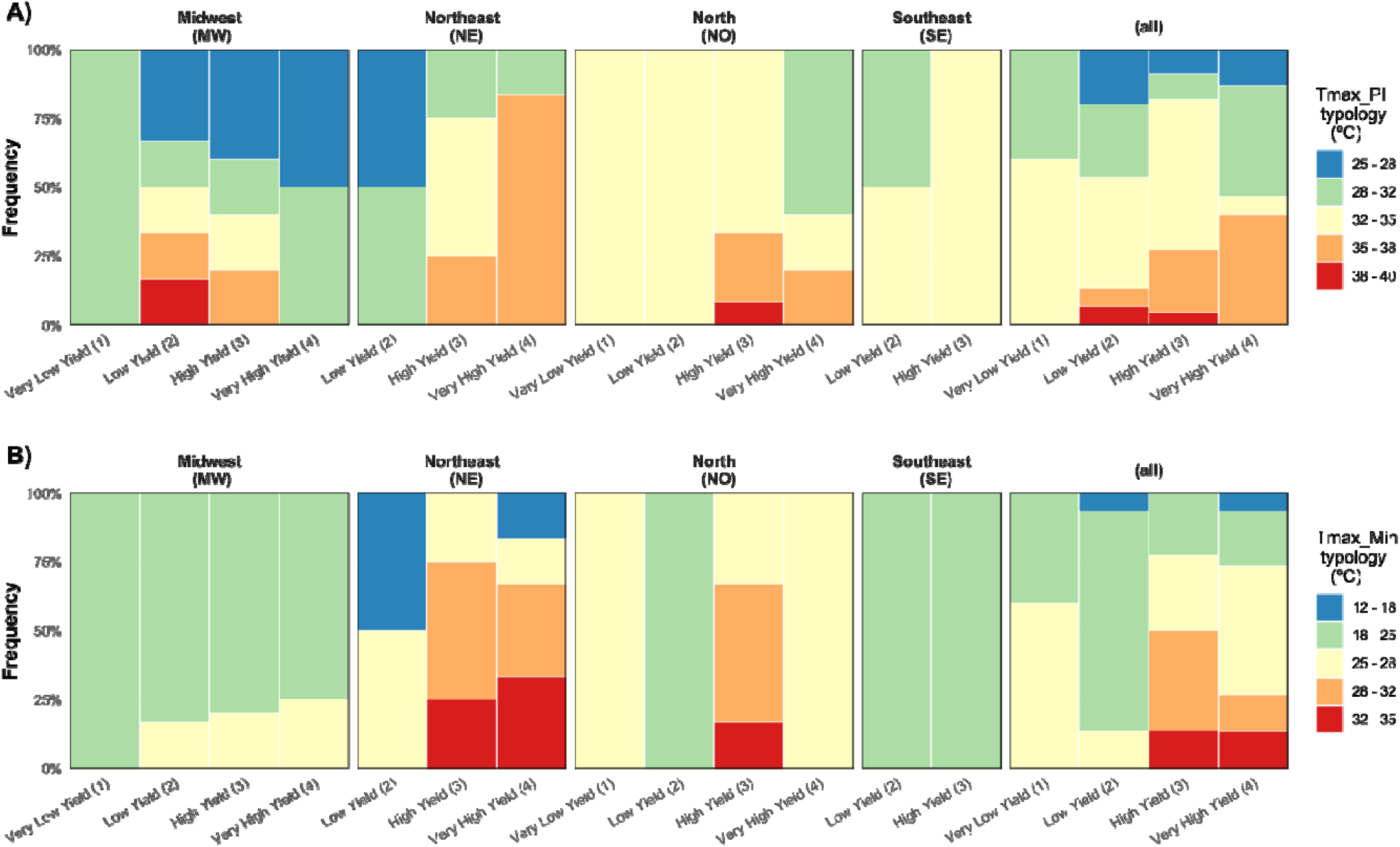
Global view of the frequency of occurrence of the main environmental covariables discriminants for genotype group “BRS Catiana” across the classifieds environments (clusters). **(A)** maximum temperature values at panicle initiation (Tmax_PI) and **(B)** Minimum value of maximum temperature (Tmax_MIN. right panel) and.

The most discriminant ECs for the “All genotypes” group were Latitude, maximum temperature value at reproductive phase (Tmax_R), and humidity at vegetative phase (Hum_V). Figure 11 shows the discrimination of Tmax_R and Hum_V across the environments (Very low yield (1), Low yield (2), High yield (3), and Very high yield (4)). Tmax_R and Hum_V ranged from 32 to 33 ºC and 60 to 65% in 50% of the occurrences at Very High (4) yield environment, respectively (Figure 11). The worst environment, very low yield (1), is characterized as having lower values of maximum temperature at reproductive phase and higher values of humidity considering 50% of the occurrences (Figure 11).

**Figure 11.**
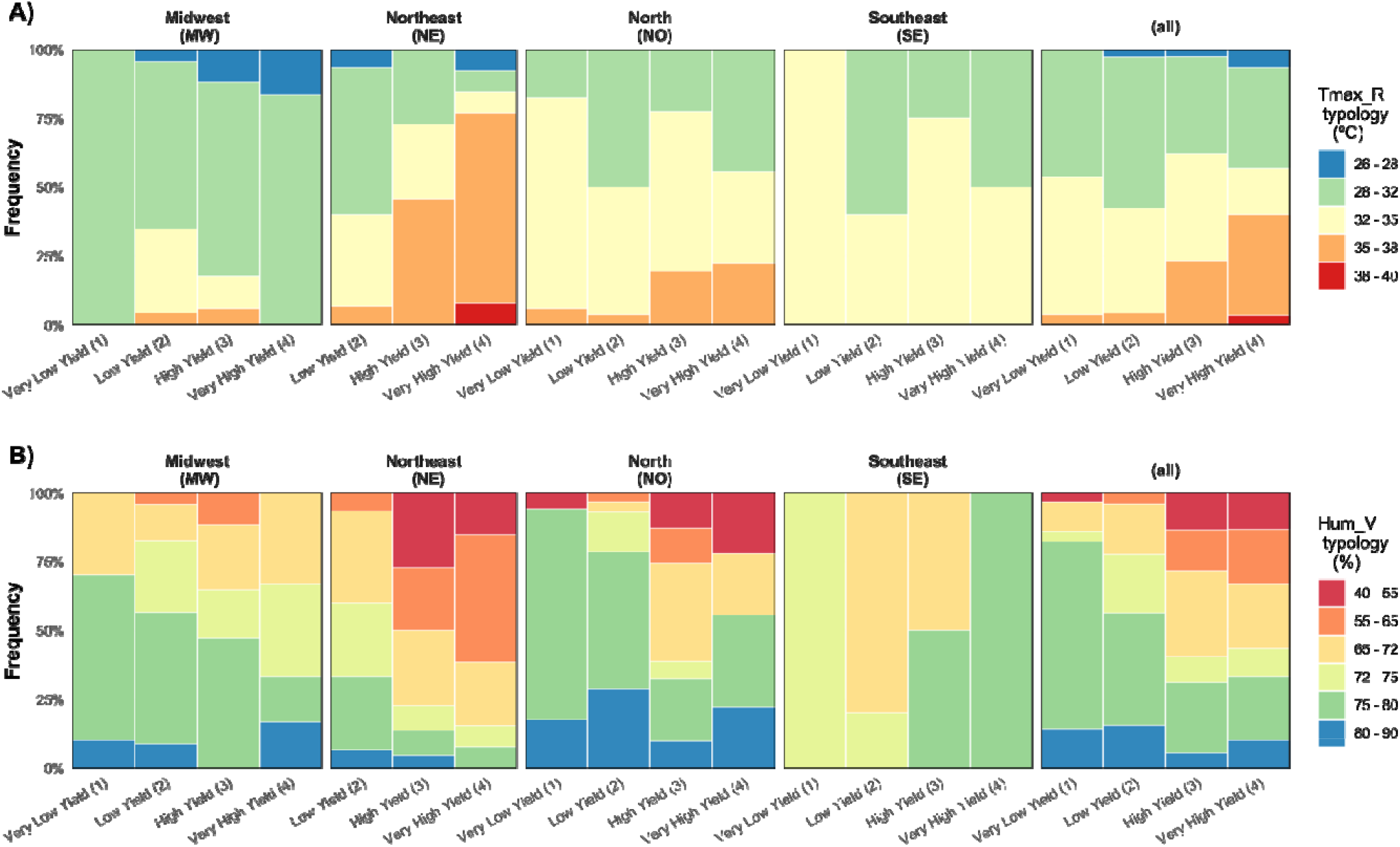
Global view of the frequency of occurrence of the main environmental covariables discriminants for genotype group “All genotypes” across the classifieds environments (clusters). **(A)** value of maximum temperature at reproductive phase (Tmax_R) and **(B)** humidity at vegetative phase (Hum_V)

### 3.6 Validation of the clusters for controlling GxE effects

Table 4 shows the results of the joint analysis of variance. In the period considered here (2011-2017), we observed that the effect of year variation (Y) is the most important source of variation for grain yield in irrigated rice in Brazil. This source represents 22% of the phenotypic variation; added with the location-year interactions, the figure increases to 34% (22%+12%). The location variation (L) is also an important driver, representing 18% of the phenotypic variation. On average, the non-genetic effects represent 51% (macro-environmental variations) and 18% (residual model effects accounting for mismodeled effects and unknown environmental sources). In practice, environmental effects are 32 times higher than broad genetic effects (G), as expected for this type of data set (advanced yield trials), with a lower genetic variability and a higher environmental diversity. These G effects represent only 9% of the phenotypic variation (which can also be interpreted as broad-sense heritability), while GE represents 22% (∼2.5x higher than G). Among GE effects, the interaction among genotypes and locations (GL, 7%) and its variation across years (GLY, 11%) has a higher contribution than the single G effect. In practice, this shows that the proportion of genetic effects in environmental and GE variations, a suggestion of the genotypic correlations, is lower (*r*_*g*_= 0.109), while the genetic correlation across locations is predominantly a crossover one (complex), with *r*_*g_l*_ = 0.334 (GL within years) and *r*_*gl_y*_ = 0.516 (GL across years). This also occurred for correlations across years, with *r*_*g_y*_ = 0.366 (GY across all years) and *r*_*gy _l*_ = 0.469 (GY within locations).

**Table 4.** Summary of the variance components and correlation parameters for each random effect of two multi-environment models with and without the yield clustering effects.

| Source | Model |  |  |  |
| --- | --- | --- | --- | --- |
|  | without clustering |  | with clustering |  |
|  | Variance Component | Explained (%) | Variance Component | Explained (%) |
| **Environment (E)** |  |  |  |  |
| Location (L) | 956.845989 | 18% | 968.205974 | 13% |
| Year (Y) | 1173.52633 | 22% | 918.306661 | 12% |
| LY | 631.542308 | 12% | 1341.455820 | 18% |
| Cluster | - | - | 1312.232270 | 18% |
| Total Environment | 2761.91462 | 51% | 4540.200720 | 61% |
| **Genotype (G)** |  |  |  |  |
| Across Clusters | 486.964429 | 9% | 450.888623 | 6% |
| Within Clusters | - | - | 375.046378 | 5% |
| Total Genotype | 486.964429 | 9% | 825.935002 | 11% |
| **G\*E Interaction** |  |  |  |  |
| GL | 360.139575 | 7% | 417.125103 | 6% |
| GY | 257.544178 | 5% | 173.011659 | 2% |
| GLY | 586.429545 | 11% | 518.667943 | 7% |
| Total G*E | 1204.113300 | 22% | 1108.804710 | 15% |
| **Residual** | 979.099452 | 18% | 922.057213 | 12% |
| **Total** | 5432.091804 | 100% | 7396.997640 | 100% |
| **Correlation Parameters** |  |  |  |  |
| $r_g$ | 0.109 | | 0.128 | |
| $r_{g_l}$ | 0.334 | | 0.606 | |
| $r_{g_y}$ | 0.366 | | 0.695 | |
| $r_{gl_y}$ | 0.591 | | 0.706 | |
| $r_{gy_l}$ | 0.559 | | 0.658 | |
| $r_{g_e}$ | 0.288 | | 0.427 | |
| **Raw proportion between the main components** |  |  |  |  |
| E : G | 5.67 |  | 5.50 |  |
| (G*E) : G | 2.47 |  | 1.34 |  |
| E : (G*E) | 2.29 |  | 4.09 |  |

The inclusion of clusters was essential for controlling them. This strategy was efficient in reducing the components related to GE (which now explains 15% of the phenotypic variation) and leveraging the environmental variation (which now explains 61 % of the phenotypic variation) mostly due to the inclusion of cluster effects (18% of the phenotypic variation). It seems that without any cluster information, both G (broad genetic variation) and GE were inflated. The broad G within clusters explains 6%, while its variation across clusters explains 5% of the natural variation from GE effects. This reflected in increasing genotypic correlations, consequently reducing the importance of crossover GE interactions, such as *r*_*g*_= 0.109 (17% of increase), *r*_*g_l*_ = 0.469 (38% of increase), *r*_*g_y*_ = 0.544 (49% of increase), *r*_*gl_y*_ = 0.706 (19% of increase), and *r*_*gy_l*_ = 0.658 (18% of increase). Considering the core of G and GE effects, the genotypic correlation across environments increased by 48%, from *r*_*g_e*_ = 0.288 (without clusters) to *r*_*g_e*_ = 0.427 (with clusters).

These results gave us a diagnosis of past GEs, but they do not really support plant breeders in the challenging task of evaluating and targeting products each year considering a different core of elite lines across multiple locations. Because of this, we detailed the study of the benefits of clustering among years and different genotype sets. Next, we investigated the benefits of clusters among years by simulating 5,000 experimental networks (different sets of genotypes and locations) across four years of real data (Figure 12).

**Figure 12.**
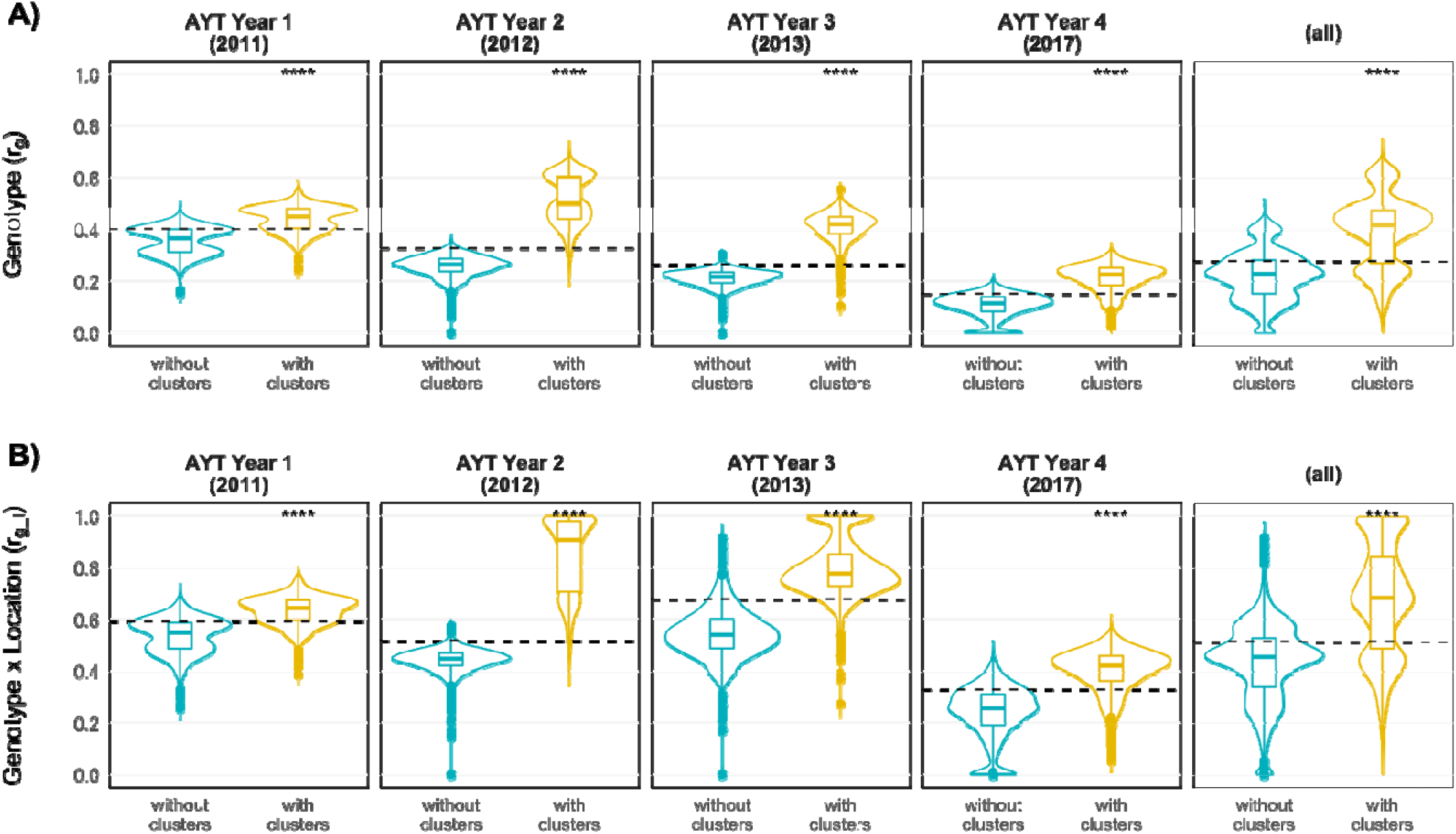
Validation of the yield clusters effectiveness in controlling genotype x location interactions (GL) across simulated experimental setups for 4 random years of real data. **(A)** Contribution of the genotypic correlations for the phenotypic variance given by the ratio of G:(G+L+GL) variance components. **(B)** Genotypic correlations across locations, given by the ratio of G:(G+GL) variance components. For this latter, values near to 1.0 represents a predominance of non-crossover (simple) GL patterns and near to 0 represents a predominance of crossover (complex) GL patterns. *** denotes highly significant differences between the model types (p < 0.001) from the non-parametric Wilcoxon test. Horizontal dashed lines represent the average value for each panel.

For all sets of years, the inclusion of clusters demonstrated a high significance (p < 0.0001, Wilcoxon test) in explaining a better capturing of G effects and controlling of GL variations, which consequently affected the estimative of genetic correlations across locations. The importance of broad genetic effect (Figure 12A), given by the average estimates of 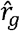 across all experimental setups and years, increased from 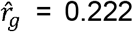 (without clusters) to 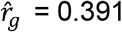 (with clusters); for some years, such as 2012, this value ranged from 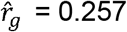 (without clusters) to 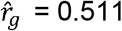 (with clusters). This means that the broad genetic effects now explain a large phenotypic variance. This also reflects in increases in genotypic correlations across locations (Figure 12B). In some years, the GL patterns went from a predominantly complex (crossover) to a non-crossover status. For example, the average values in the AYT sets of 2012 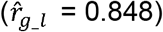 and 2013 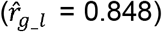 suggest a high predominance of simple GL patterns, which may facilitate strategies of product development and cultivar recommendations. As the ML approach focuses on predicting environmental variations, it is reasonable to assume that all other correlations may also be predicted using additional information (in our case, environmental covariables), as we discuss below.

## 4. DISCUSSION

The sustainability of rice production depends on the development of new rice cultivars with high yields and stable performance across diverse environments (Donoso-Ñanculao et al., 2016). Therefore, one of the major challenges in plant breeding is the differential response of genotypes in different environments, known as GE interaction. Finding lines with predictable GE across seasons (GY) has become an increasingly pressing issue due to the highly variable and unpredictable weather conditions in current and future climate change scenarios (Monteverde et al., 2018). Multi-environment studies play a critical role in this matter. Several approaches for modeling GE have been proposed in the literature (Heslot et al., 2014; Donoso-Ñanculao et al., 2016; Monteverde et al., 2018); all confirm the relative benefits of using multi-environment models in comparison with single-environment analyses.

GE variation represents an opportunity for selection for adaptation, as it shows that some genotypes perform better in some specific growing conditions than others. However, in practical terms, for plant breeding the GE represents a bottleneck for a successful release of a new cultivar, especially at late stages of breeding and even at post-breeding stages, such as cultivar recommendation and seed production. The main issue plant breeders face is the lack of trust that the environmental quality of the experimental network really reflects the environmental quality (and diversity) of the main target population of environments (TPE). The TPE represents the expectation of growing conditions to which a certain product (cultivar) was designed and recommended. However, a main issue emerges: what connects these breeders’ expectations to the reality of farm fields? The answer is (1) agronomic expertise, mostly related to knowledge about the environmental background of a certain region, which involves not only regional/geographic variations in edaphoclimatic factors but also knowledge about some diseases in certain regions and how seasonal factors tend to occur and be deal with (e.g., agronomic practices to deal with drought-stress); (2) a data-driven approach mostly relating yield variations, such as empirical characterizations, using observed data for a given trait or environmental data (envirotyping-based characterization, Crespo-Herrera et al., 2021), which may involve the use of simulated yield levels and crop growth models (Chenu et al., 2011; Heinemann et al., 2008, 2015, 2019) or even farm yields from public databases (Heinemann et al., 2011).

In our study, we introduce the use of an unsupervised pattern recognition algorithm to conciliate the breeders’ expertise with the data-driven empirical characterization of the culture region. In this first step, empirical data are used to reveal the “putative clusters of yield levels.” Next, we introduce the use of a supervised machine learning approach capable to link the empirical clusters to actual envirotyping data. An interesting point of this approach is the fact that it may be used for a core of genotypes, such as the “all genotypes” group evaluated in this study, or even for cultivar-specific variations.

Our approach was efficient in recognizing the empirical patterns related to genotypic variation, climatic (seasonal) factors, and repeatable geographic differences across years for grain yield in irrigated rice. From the advanced yield trials (AYT) of irrigated rice breeding programs in Brazil, we could identify that the proportion between broad genotypic effects, environmental variation, and genotype by environment interactions for grain yield is approximately 1.00 (G):5.67 (E):2.47 (GE), which is almost 1:6:2. This suggests that for any location the GE variation is at least two times higher than the G variation, which consequently may hinder product development for this huge, broad culture region. However, after the identification of clusters of yield levels, it seems that the GE pattern is now predominantly a non-crossover one (simple). After including the cluster information in GE models, the relation between G:E:GE decreases to 1.00 (G):5.50 (E):1.34 (GE), which is almost 1:5:1. In addition, after our simulation, we could validate these clusters as homogenous mega-environments, which may be used in the future to subdivide the current TPE into small TPEs with specific product design goals.

Donoso-Ñanculao et al. (2016) estimated the components of variance for grain yield evaluations in three locations for three years in Chile using a mixed model. The results were similar as those of the ANOVA analysis here. Genotype was the most important variation source (25%). The importance of year (8.7%) was intermediate; the significance of location was low; and there was a high G×Y interaction. The important contribution of genotype and the low contribution of location may be explained by the small size of the Chilean rice zone. In our study, there was a low contribution of genotype to the variance component probably because of the high variance in climate due to the broad Brazilian rice area. Published rice multi-environment studies generally report a high contribution of environment or G×E interaction (Samonte et al., 2005; Camargo-Buitrago et al., 2011).

One of the goals of product design, for instance, is the identification of which environmental factors should drive each yield level, instead of using environmental factors to define possible clusters of yield levels (Crespo-Herrera et al., 2021). In addition, the use of envirotyping to predict actual yield clusters may be an important source of information related to unrevealing the causes for such a low yield (low yield groups). It should highlight possible gaps in product design and development since the early stages of a breeding program. This allows the implementation of new strategies that account for envirotyping uses as a tool for harnessing breeding and agronomic practices that mitigate the lack of adaptation to a certain region (Cooper and Messina, 2021). This is an additional benefit of our data-driven strategy developed, which creates the possibility of measuring the impacts of environmental covariables for specific genotype groups and may involve a core of genotypes (e.g., “All genotypes”) or cultivar-specific data (e.g., IRGA 424 and BRS Catiana).

The usefulness of pinpointing these impacts aids producers in developing adaptation strategies based on varietal selection, which seems to represent a major contribution to irrigated rice breeding efforts that seek to develop better varieties adapted to tropical regions. Thus, we may enumerate two uses of this approach involving the use of envirotyping data in yield clusters: (Strategy 1) understanding specific patterns of adaptation of genotypes, especially for pinpointing the main environmental typologies driving reaction norms; and (Strategy 2) understanding the expected yield levels for a certain region and considering a certain germplasm. Below we detail each of this uses.

Our results showed that temperature is an issue for irrigated rice in tropical regions. Maximum temperature at filling grain, flowering, and reproductive phases are ECs discriminants for cultivars such as IRGA 424, which seems not adapted to the climatic conditions of the NE region. Pereira et al. (2011) found that although IRGA 424 does not present agronomic or market limitations for cultivation in the NE region, it has ranked below BRS Catiana and other cultivars in environments that are more or less conducive to higher yields.

High temperature stress is one of the major concerns. It is considered as a result of climate change. This is a possible example of using the Strategy 1. Another use of Strategy 1 is to connect agronomic practices, such as adjustment of planting dates, to the current framework for cultivar testing.

In Scenario 2, from an agrometeorological point of view, our approach was also efficient in predicting yield levels, which may leverage decision-making about product targeting (recommendation of cultivars for certain regions and planting dates). In addition, it may be a powerful tool supporting other post-breeding operations, such as selecting the locations with the highest probability of achieving higher yield levels, which in turn may guide the implementation of seed productions less prone to yield losses. For example, our study showed that the regions Northeast (PI and States) and Southeast (SP State) could be the main places for seed production. They are defined by high-quality environments with potential to achieve higher yield levels. The strategy can also be used for product targeting, in which a single genotype (e.g., BRS Catiana or IRGA 424) can be used to fit the model and then explore the differential reaction norms in terms of differential typologies.

The ability of models to predict GxY is crucial for crop improvement in current and future climate change scenarios. This is likely to be especially beneficial for rice breeding programs that focus on high yield and quality, given that both components are highly affected by environmental conditions (Cameron and Wang, 2005; Chen et al., 2012; Monteverde et al., 2018).

With large-scale envirotyping, in combination with known genotypic information, environments for crops may be optimized and phenotypic performance under specific environments may be largely predicted, thus improving cost-benefit ratio for precision breeding and crop production (Xu, 2016).

## 5. CONCLUSION

We conclude that the data-driven approach is efficient in understanding the empirical patterns of yield variation and translating it into predictable patterns of environmental quality. This tool proves its usefulness in understanding the climatic and geographic drivers of GE for irrigated rice in Brazil, highlighting how seasonal variations in those environmental factors contribute to differentiate the quality of irrigated rice fields across years and diverse regions in Brazil. In practical terms, this approach delivers a useful information for designing clusters of homogenous environments, which enhances the ability of GE models used for analyzing nationwide data aiming cultivar targeting. However, it also provides reliable information about the current yield levels of certain Brazilian regions. Below we enumerate the benefits of using our approach:

**Benefit 1:** Ease the use of long-term historical yield data to group field trials based on identified clusters; if analyzed across years, it can reveal the most prone yield levels expected for a given location at a certain planting date. This may make the zoning of cultivars easier and guide practices that address agronomic climatic risks directly from the breeding field trials to the actual farm fields.

**Benefit 2:** Support the definition of expected yield levels for different levels of analysis, since the study of certain regions based on past yield trial data and their associations with geographic and climatic factors, which mostly represents static and seasonal patterns of environmental quality conditions, and including the study of a whole germplasm for a given region (across regions) or even the prospection of how the environmental variables affect the adaptability of a given cultivar.

**Benefit 3:** Provide information for different stages of varietal research, such as (a) product design, by understanding the limits of yield level for a certain target region; (b) product development, by accelerating the process of cultivar recommendation and agronomic practices definitions; and (c) at post-breeding stages of seed production, by supporting the selection of locations most prone to achieve higher yields, which represents the optimization of resources and an increase in security as for not selecting production fields prone to lower yield levels.

## ACKNOWLEDGMENTS

AB Heinemann acknowledges support from “Fundação de Amparo à Pesquisa do Estado de Goiás” (FAPEG -PRONEM/FAPEG/CNPq) and “Conselho Nacional de Desenvolvimento Científico e Tecnológico” (CNPq –Edital Universal –Processo -408025/2018-2

