## Supplementary for "Data-driven machine learning for pattern recognition supports environmental quality prediction for irrigated rice in Brazil"

**RUN TITLE**: Machine learning-based enviromics prediction of yield levels for rainfed rice in Brazil

**Highlights**

- A nationwide environmental characterization and its relation to the genotype by environment interaction (GE) for grain yield of rainfed rice growing regions in Brazil.
- A data-driven approach capable to identifying clusters of yield levels and a machine learning approach to relate those clusters with environmental typologies.
- Unrevealed geographic and climatic causes of environmental quality for a group of genotypes or cultivar-specific predictions.
- The strategy benefits diverse stages of breeding (multiple environmental trial analysis) and post-breeding (selection of fields for seed production) as an alternative approach to reduce costs and support decisions on cultivar planting locations.

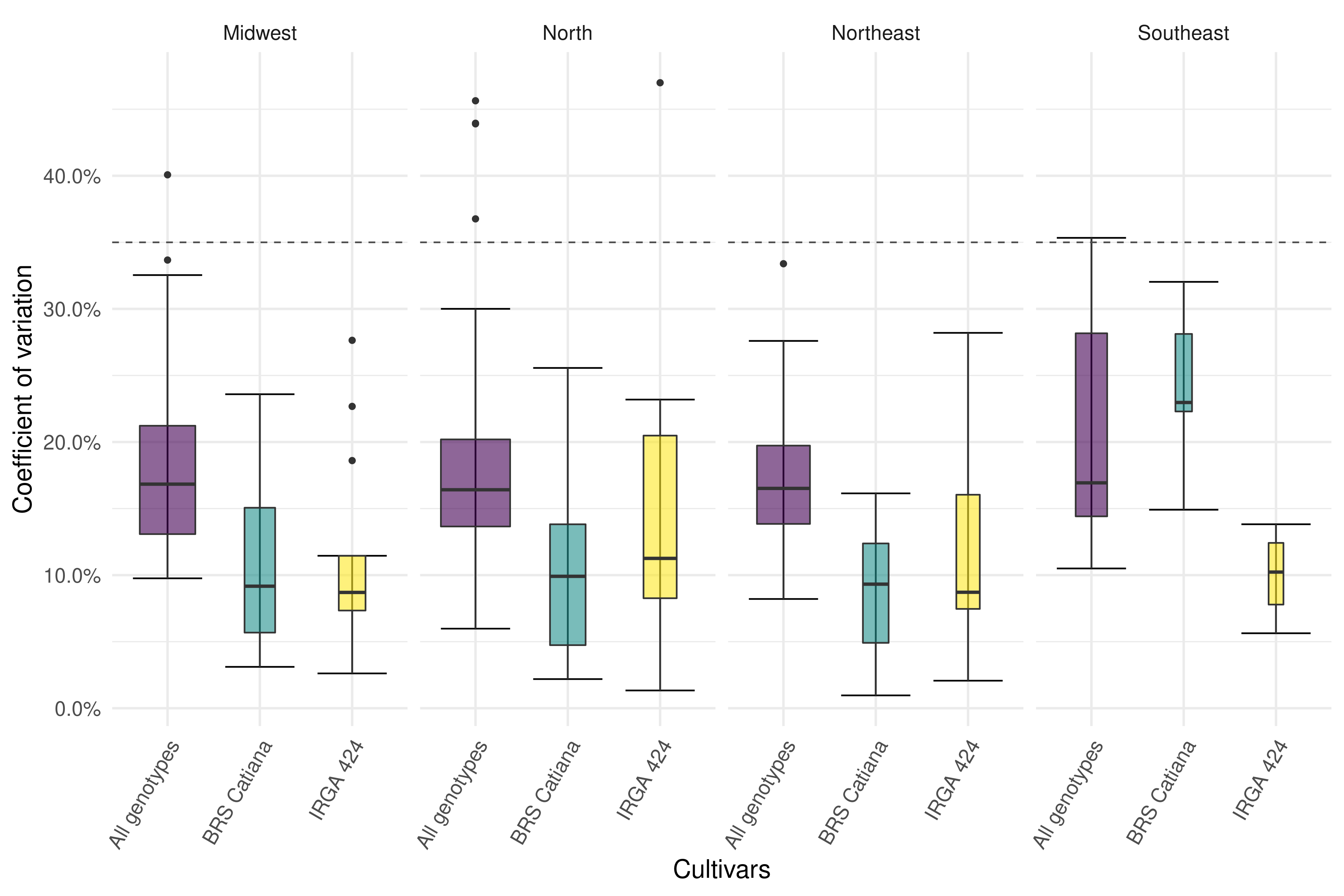

Figure S1. Coefficient of variation of irrrigated yields trials across megaregions (Midwest, North, Northeast and Southeast, top panel).

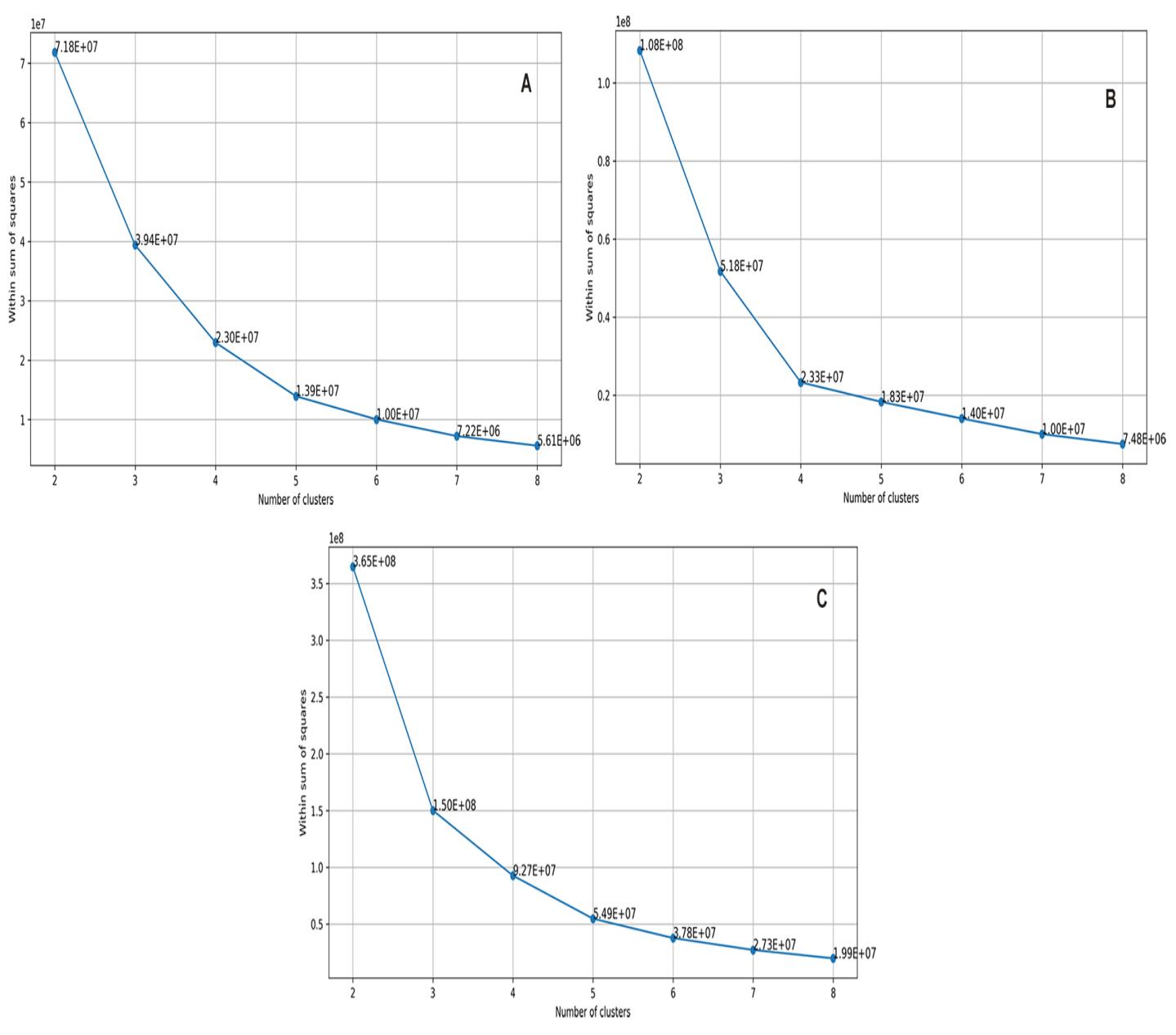

Figure S2. Elbow method based on the within cluster sums of squares for genotype groups A) IRGA 424, B) BRS Catiana and C) All genotypes

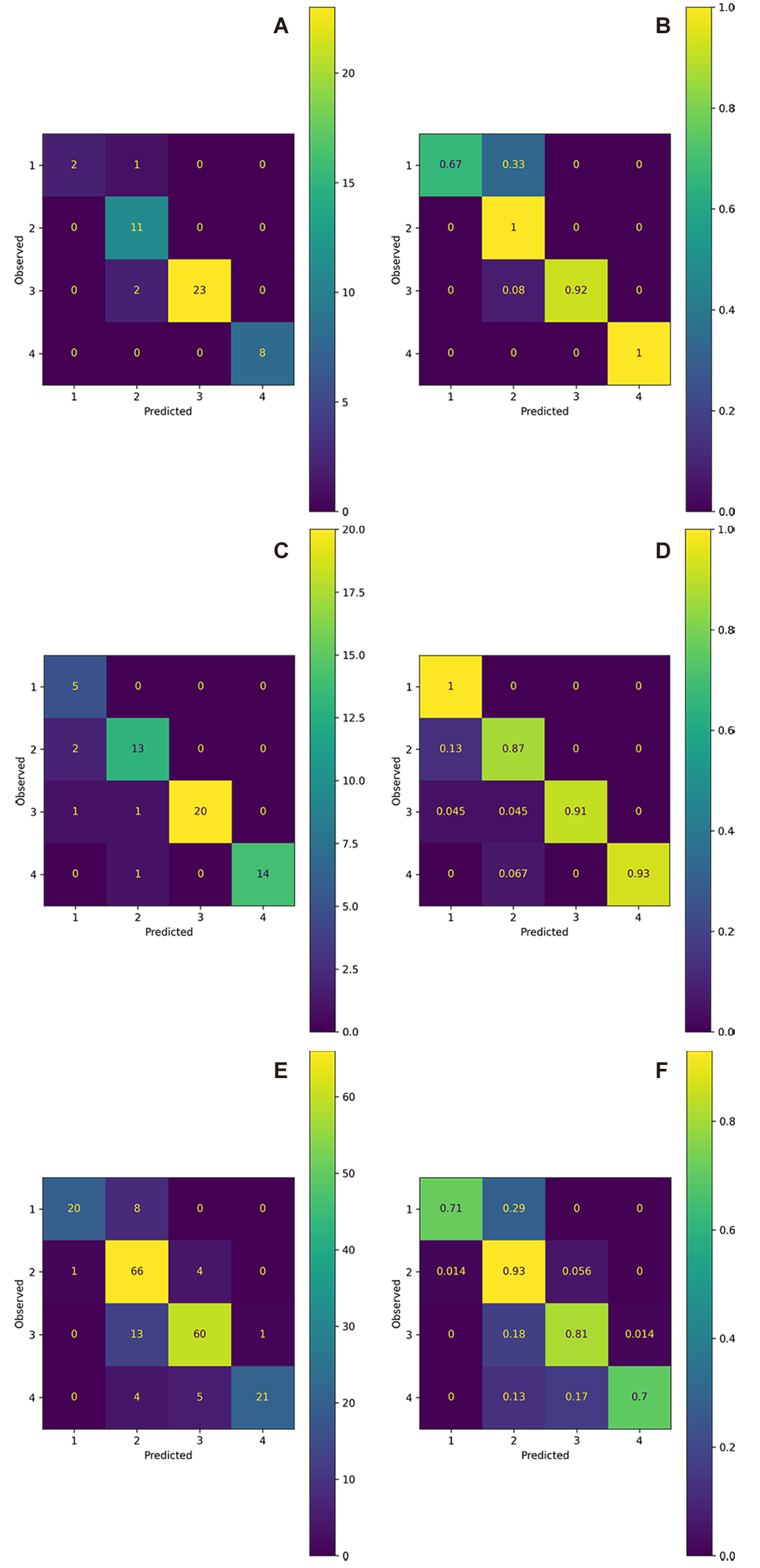

Figure S3. Matrix of confusion showing the decision tree classification models precision for the genotypes groups A) and B) IRGA 424, C) and D) BRS Catiana and E) and F) All genotypes. A), C) and E) shows the true positive (TP), the number of cases correctly identified as positive in the diagonal. B), D) and F) shows the percentage of the true positive (TP) in the diagonal. Outside of the diagonal are the false positive (FP), the number of cases incorrectly identified as positive.
